# Insights into actin polymerization and nucleation using a coarse grained model

**DOI:** 10.1101/715383

**Authors:** Brandon G. Horan, Aaron R. Hall, Dimitrios Vavylonis

## Abstract

We studied actin filament polymerization and nucleation with molecular dynamics simulations and a previously established coarse-grained model having each residue represented by a single interaction site located at the C*_α_* atom. We approximate each actin protein as a fully or partially rigid unit to identify the equilibrium structural ensemble of interprotein complexes. Monomers in the F-actin configuration bound to both barbed and pointed ends of a short F-actin filament at the anticipated locations for polymerization. Binding at both ends occurred with similar affinity. Contacts between residues of the incoming subunit and the short filament were consistent with expectation from models based on crystallography, X-ray diffraction and cryo-electron microscopy. Binding at the barbed and pointed end also occurred at an angle with respect to the polymerizable bound structure, and the angle range depended on the flexibility of the D-loop. Additional barbed end bound states were seen when the incoming subunit was in the G-actin form. Consistent with an activation barrier for pointed end polymerization, G-actin did not bind at an F-actin pointed end. In all cases, binding at the barbed end also occurred in a configuration similar to the antiparallel (lower) dimer. Individual monomers bound each other in a short-pitch helix complex in addition to other configurations, with several of them apparently non-productive for polymerization. Simulations with multiple monomers in the F-actin form show assembly into filaments as well as transient aggregates at the barbed end. We discuss the implications of these observations on the kinetic pathway of actin filament nucleation and polymerization and possibilities for future improvements of the coarse-grained model.

**SIGNIFICANCE:** Control of actin filament nucleation and elongation has crucial importance to cellular life. We show that coarse-grained molecular dynamics simulations are a powerful tool which can gauge involved mechanisms at reasonable computational cost, while retaining essential features of the fully atomic, yet less computationally tractable, system. Using a knowledge-based potential demonstrates the power of these methods for explaining and reproducing polymerization. Intermediate actin complexes identified in the simulations may play critical roles in the kinetic pathways of actin polymerization which may have been difficult to observe in prior experiments. These methods have been sparsely applied to the actin system, yet have potential to answer many important questions in the field.

## 1 INTRODUCTION

The polymerization of actin proteins into filaments is of fundamental importance in basic cellular functions such as cell migration, cytokinesis, and neuron growth (1). A long history of thermodynamic, kinetic and structural studies of actin established that conversion of actin monomers (G-actin) into filaments (F-actin) involves the creation of a nucleus of 3-4 subunits that grows by polymerization above a critical monomer concentration ∼ 0.1 *µM* (under typical polymerization buffer conditions) (2–4). The free energy of hydrolysis of monomer-bound ATP, which occurs following polymerization and associated with a conformational change of the actin molecule from G to F form, maintains a treadmilling steady state with net polymerization (depolymerization) at the barbed (pointed) end, even in the absence of cellular co-factors that further accelerate turnover by several times in cells (3, 5, 6). ATP-actin polymerizes about ten times faster at the barbed end (rate constant *k*^+^ ≈ 10 µ*M s*) as compared to the pointed end (7). Polymerization at the pointed end is thought to involve conformational changes (8, 9) that require overcoming an activation barrier, as evidenced by the dependence of polymerization rate constant on viscoscity (10).

Models of actin filament elongation typically assume polymerization and depolymerization occurs monomer by monomer, in a process that can be described as a random walk and simple chemical kinetics (11–13). However, the intermediate structures involved in polymerization have not been elucidated by modeling methods in molecular details yet. It is known that actin monomers may interact with each other in ways that differ from the native contacts of the actin filament double helix. For example, actin monomers can also bind in an antiparallel manner (the lower dimer configuration) (14–18) and complexes unproductive for polymerization have been observed in the early stages of nucleation (19). Measured filament length fluctuations at steady state in vitro (slightly above the critical concentration of the barbed end) were larger than those expected by a monomer by monomer association and dissociation process (20). This observation lead to the suggestion that the polymerization unit may be an average of monomers, oligomers and fragments (20). Hydrolysis of ATP bound to actin would contribute to length fluctuations, however kinetic models suggest hydrolysis-induced enhanced length fluctuations occur slightly below the barbed end critical concentration rather than slightly above (12, 13).

Several modeling studies have examined structural and conformational changes occurring within the actin filament, using both atomistic and coarse-grained representations (21–27), however not much is known about the interaction ensemble involved in polymerization and nucleation. This is a crucial aspect of the actin system as its key regulators are nucleation- and elongation-promoting factors, such as formins, Arp2/3 complex and Ena/VASP proteins, which form large complexes whose aim is to bring together actin monomers and guide them to the barbed end much more rapidly than they would otherwise on their own (3, 5).

In pioneering works, Sept et al. (28, 29) used binding free energy calculations and Brownian dynamics simulations with atomistic level of detail to evaluate both equilibrium and kinetic constants of polymerization and dimer/trimer formation. However these authors assumed knowledge of possible geometric interactions between actin proteins and did not consider the full equilibrium ensemble. More recently, Ohnuki et al. (30) examined the electrostatic energy of subunit addition to the barbed end using all-atom actin filament structures via energy minimization, without however exploring the full configurational space.

In this work, we utilize molecular dynamics (MD) simulations to simulate the formation of multi-subunit actin complexes using the C_*α*_ level model of Kim and Hummer (KH) which was designed and tested for coarse-grained simulations of multiprotein complex formation (31, 32). We recently showed this model was suitable to describe the Formin Homology 1 (FH1) domain, an intrinsically disordered region (33). The model was also able to simulate the delivery and binding of profilin-actin to the barbed end (33), hence it may have great potential to capture nm and sub-nm features important in future modeling of cytoskeletal filament polymerization and nucleation and their regulation by multi-protein complexes. This manuscript has thus two goals. First, to take a closer look at how the KH model describes known features of pure actin polymerization as well as the more elusive aspects of pure actin nucleation. Secondly, to use the model, within its resolution limits, to study mechanistic aspects of polymerization and nucleation that other prior models have not yet explored.

We describe the simulated structural ensemble of an actin monomer binding to an actin filament seed (a rigid dimer) and how it depends on the conformation of the incoming monomer. We consistently observe an antiparallel dimer complex forming at the barbed end, as well as other complexes that have not been previously identified experimentally. Individual monomers bound each other in multiple configurations, with several of them apparently not favoring the continuation of polymerization. Simulations with multiple monomers in the F-actin form show assembly into filaments as well as transient aggregates at the barbed end. We discuss the implications of our results for the kinetic pathway of actin filament nucleation and polymerization and on the development of improved coarse-grained models.

## 2 METHODS

We utilize molecular dynamics simulations to simulate the formation of multi-subunit actin complexes. We use the model of Kim and Hummer (KH) (31) where each amino acid is represented with a single bead at the location of the alpha carbon. Specifically, in the KH model, nonbonded pairwise interactions are modeled by a Lennard-Jones (LJ) type potential, which is either attractive or repulsive, based on the Miyazawa-Jerningan pairwise interaction matrix (34). Debye-Hückel electrostatics at ∼ 100 mM salt apply for all pairs of charged residues, implemented as a screening length of 10 Å and uniform dielectric constant 80. In simulations with flexible domains, the beads are connected along a chain by harmonic springs with equilibrium length 3.81Å and spring constant 756 kcal/(mol Å^2^). For simplicity, other terms in the potential energy function due to angular or dihedral constraints are not included when considering flexible domains.

The KH model was developed in different versions (Models A-F) to account for solvent accessibility. We performed simulations with Model A, in which all residues are weighted equally irrespective of their distance from the protein surface. This model does not involve a solvent accessibility calculation at every time step and it is thus much less computationally costly for simulations with flexible domains. Model A was somewhat sensitive to the LJ interaction cutoff when applied at 3 versus 4 *σ* of the corresponding potential, with the 4 *σ* cutoff model giving *K_d_* values that were lower by up to two orders of magnitude and favoring the antiparallel binding mode over other modes (the antiparallel structure being the the most compact structure and thus having the strongest interactions among interior residues). For most results below we thus used Model A with a 3 *σ* cutoff. For better sampling of bound states we used a reduced temperature (197 K). We tested that while the reduced temperature influenced the overall binding strength by several orders of magnitude, it did not influence the ratio of probabilities of bound states by more than a factor of 5.

MD simulations were performed in the canonical ensemble using LAMMPS (35, 36), see Supplemental Text for more details. Except where noted, simulations were conducted using the replica exchange molecular dynamics (REMD, parallel tempering) method, which allows for probing of the equilibrium ensemble with reasonable time and computational cost (37).

To calculate the dissociation constant between species with multiple binding modes *i*, we used the law of mass action in simulations that contained one protein of each kind and varying volumes:

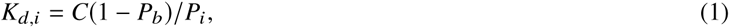

where *C* is the concentration of one protein in the simulation box and *P_i_* is the probability to be in a given binding mode. *P_b_* is the total probability of the two species being bound, defined as all frames which have negative interaction energy between the two species (the population of configurations with energies around 0 were either identically 0 or slightly positive).

In all simulations we did not include the nucleotide bound to actin, which is not in a position to make direct contacts with residues of neighboring subunits. The electrostatic energy between the ADP and a dimer with bound ADP would be less than 3% of the calculated energies of bound complexes, within the KH model used. Experimentally, nucleotide-free actin establishes similar intersubunit contacts along the long-pitch helix to ADP-actin (38). We also do not consider the effects of divalent ions stabilizing both long- and short-pitch helix contacts (39, 40).

The residues missing in the G-actin crystal structure (41) were added using the MODELLER software (42). For simulations with flexibility, the same residues that were missing from the G-actin crystal structure we used were treated as flexible: the D-loop (r40-51) and the termini (r1-5, r372-375). For the simulations with partially flexible monomer binding to a dimer, the two termini of each dimer subunit (located at and close to the barbed end) were flexible, but the D-loops of both (located at the pointed end) were kept rigid in the Oda et al. (43) conformation for reasons of computational efficiency.

We used distance root mean square (dRMS (31)) as measure to quantify similarity to a reference bound structure. If dRMS is sufficiently low, as described below, then a simulation frame is considered successfully bound in the reference structure. (31). This is performed by calculating the average absolute deviation in distances between alpha carbon residues from the test structure and residues from reference structure near the binding interface. We use a cutoff of 10Å in the reference structure to determine which pairs of residues to calculate distances between.

We also used rotation and translation with respect to the reference F-actin structure of Oda et al. (43) to quantify bound states of monomers to a minimal F-actin dimer filament seed. We calculated a rigid body transformation, *x*′ = *Rx* + *T* (where *x* and *x*′ are the 3D vectors), to transform the coordinates of the monomer to those of a monomer bound at the barbed end in the Oda et al. position. For simulations with a G-actin monomer, we replaced the F-Oda monomer with a G-actin monomer in the reference structures by alignment. We then diagonalized the rotation matrix *R* to extract a single angle of rotation by taking the positive imaginary eigenvalue. The components of *R* and *T* were calculated using four reference residues of the incoming monomer chosen near the corners of the monomer.

To classify and discriminate different binding modes of monomers to F-actin dimer, we created two-dimensional distributions showing the translation and rotation of the monomer with respect to the predicted barbed end binding location in the Oda et al. structure as described above. We also used two-dimensional dRMS plots from the predicted barbed and pointed end locations, which helped resolve the degeneracy of high dRMS values with respect to either location. For the monomer binding to another monomer simulations, we used the Oda et al. short-pitch helix and long-pitch helix as our reference structures for two-dimensional distributions.

For each simulation, we calculated interaction energy, dRMS with respect to one or two reference structures, and rotation angle and translation magnitude (Fig. S1). Unless otherwise stated, probability of a given binding mode was defined using the two-dimensional distributions in the following manner. We first identified all of the maximal bins of the distribution which contained at least 100 data points in BE/PE dRMS plots (Fig. S1B) binned at 0.1 Å. Then we identified a region around the maximal bins in any direction for which all points in the region had a probability of at least 1.8% of the probability of the corresponding maximal bin (i.e. treating the distributions as if they were Gaussian distributions, uncorrelated in either direction, and using a cutoff of two standard deviations in either direction to determine whether a point should be in the region). Then, if regions were found to be overlapping, they were subsequently divided at the local minimal bins between the regions, and the probability of the minimal bins was equally divided by the number of regions they shared. Occasionally multiple regions were identified that were too close to be classified as distinguishable regions (dRMS < 3 Å). These regions were then combined as a single region. Only regions with total probability ≥ 1% based on this definition were labeled. As needed, convergence of simulations was determined by ensuring the relative error of the maximum in the probability bins that define bound states in the two-dimensional distributions was sufficiently low (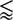10%) across multiple independent simulations, i.e. we check that the probability to visit each binding mode has sufficiently converged across different simulations.

## 3 RESULTS

To study the molecular interactions during actin polymerization and nucleation, we performed coarse-grained MD simulations. We used the knowledge-based potential of Kim and Hummer (31), developed for multiprotein complex formation. This model has been successfully applied to predict the various conformation of bound structures, binding affinities, and protein encounter complexes (31, 32). We recently showed this model was suitable to describe the Formin Homology 1 (FH1) and the transfer of profilin-actin bound to the FH1 domain to the barbed end (BE) of an actin filament (33, 44). To achieve fast convergence to thermodynamic equilibrium, we used the replica exchange molecular dynamics (REMD) method (37) and assumed actin monomers have rigid shape (or partly rigid, as described below).

The model of Oda et al. (43) for F-ADP-actin was used to define the structure of F-actin in our simulations. This model, which is based on X-ray diffraction data, has been updated by models using cryo-EM, the resolution of which has been increasing study after study (9, 45–50). We did not explore the differences of these structures to Oda et al. that should be small for the purposes of this work. In the Oda et al. model, contacts stabilizing the actin filament form along the long-pitch helix direction (also known as longitudinal or intrastrand direction) and short-pitch helix direction (diagonal or interstrand), see Fig. 1A. The contacts along the long-pitch direction are estimated to be stronger (9, 28, 49, 51, 52). These involve contacts between subdomain 4 of the subunit towards the BE with subdomain 3 of the subunit towards the pointed end (PE). The D-loop, a flexible region in subdomain 2 (53–55) of the subunit towards the BE is involved in contacts with the hydrophobic groove between subdomains 1 and 3 of the the subunit towards the PE. Binding along the short-pitch helix largely involves contacts between subdomains 3 and 4.

**Figure 1:**
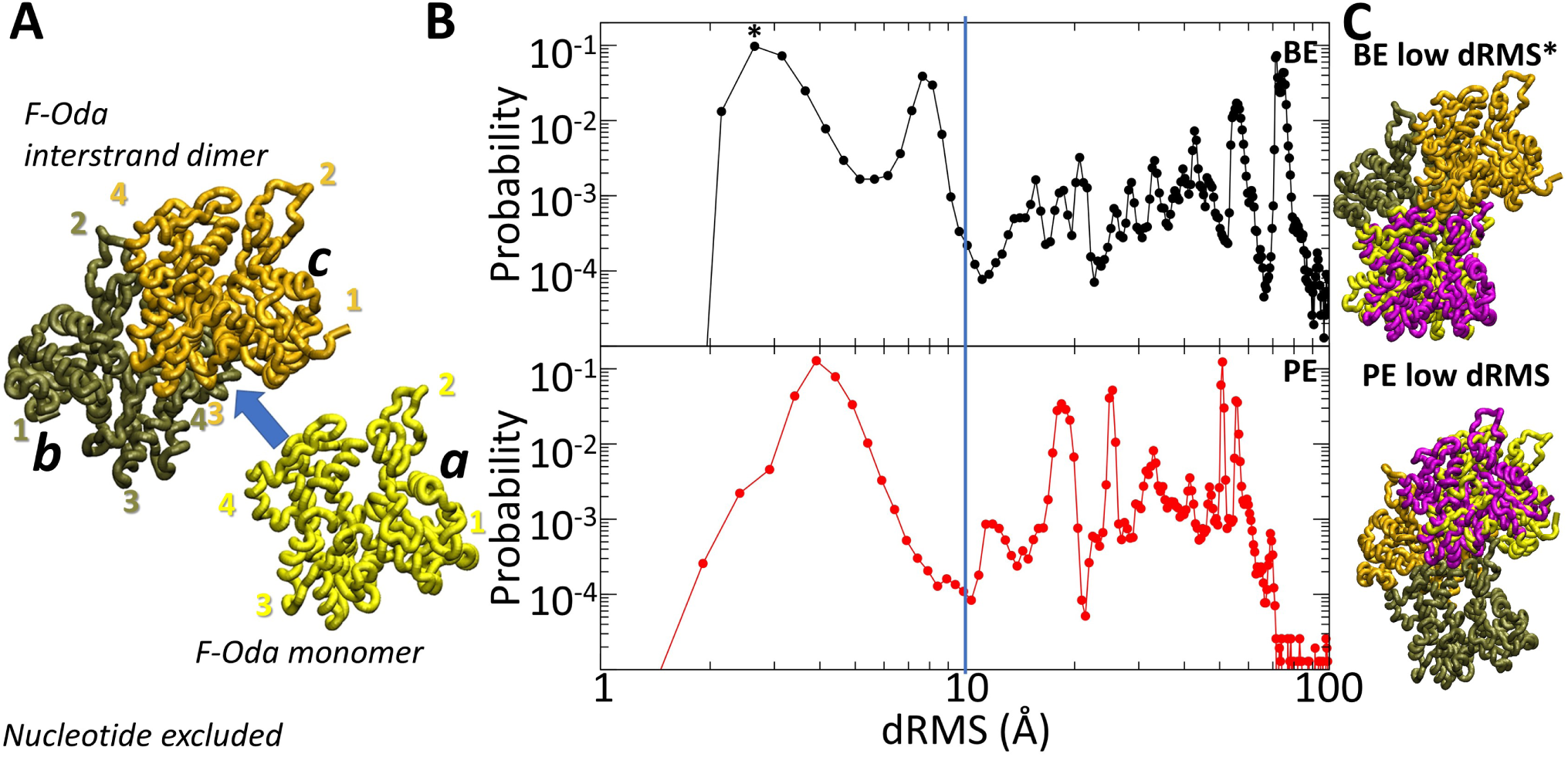
Coarse-grained KH model predicts actin monomer association to barbed and pointed ends. (A) Cartoon depicting the simulation of a monomer (subunit *a*) in binding equilibrium with a dimer of subunits *b* and *c* rigidly connected to one another as a minimal F-actin filament exhibiting barbed (bottom) and pointed (top) ends. This figure shows results for subunits in the rigid F-Oda configuration. The four subdomains of each protein are labeled. (B) Distribution of dRMS of subunit *a* with respect to the BE or PE position in the structure by Oda et al. (43). Data are for Model A with 3*σ* cutoff at 197 K in box of size 79 nm corresponding to concentration 3.3 *µM*. (C) Images comparing the structures at the lowest dRMS peaks at BE and PE (subunit *a* in yellow) to the Oda et al. structure (pink).

### Simulations of rigid monomers binding to the the barbed and pointed end

To get a baseline reference for how well the model predicts the association of monomers to the filament ends, we performed simulations of a rigid short-pitch helix (interstrand) dimer (“F-Oda dimer”) interacting with a rigid “F-Oda” monomer, namely a monomer with the same structure as an F-actin subunit in the Oda et al. model (Fig. 1A). The two subunits of the dimer where moved together as a single rigid body. This setup is the simplest possible setup which allows all filamentous contacts between the dimer and the monomer to form. It allows us to test how well the KH model can capture actin interactions but does not yet consider the complexities associated with the conformational transition from a twisted conformation (G-actin) of the incoming monomer in the bulk to a flattened conformation in the filament (F-actin) (5).

Without making any assumptions about the types of complexes that may form in the simulation, we examined the probability of the monomer to bind to the BE or PE. To quantify this probability, we used the dRMS distance of the F-Oda monomer with respect to the BE/PE reference structure in the Oda et al. model. Structures with dRMS < 10 Å to either the BE or PE were considered consistent with the Oda et al. model for a monomer in the interior of the filament.

We found that the monomer did indeed bind to the BE, very close to location anticipated in the Oda et al. model (peaks at 3 and 8 Å in Fig. 1B,C). Since binding was weak at high temperatures (Fig. S2), and since the binding ensemble and relative binding probabilities did not change significantly with temperature, we used one of the lower temperatures of the replica exchange simulations (*T* = 197 K) to report binding probabilities (temperature in these simulations changes the relative magnitude of rigid body translational and rotational entropy loss upon binding with respect to the temperature-independent KH interaction strength gain). At this reduced temperature, the calculated dissociation equilibrium constant was 0.15 *µ*M (Table S1, Fig. S2). The dissociation equilibrium constant at 300 K was in the range of 10^4^ *µM* (Fig. S2A), much higher than the critical concentrations of ATP-actin (∼ 0.15 *µ*M (56)) and ADP-actin (~1*µ*M (39)) at 100 mM salt assumed in the Debye-Hückel expression. This difference is likely due to interactions enhancing binding that are beyond the resolution of the KH model (see Discussion section).

We also found that the F-Oda monomer bound to the PE, very close to the anticipated location, and with almost identical affinity as the BE (peak at 4 Å in Fig. 1C and Fig. S2). Our approach is thus self-consinstent: if all monomers in a trimer are flat and close to the F-Oda configuration, its free energy would not depend on whether binding of the third subunit occurs at the BE or PE. One would thus expect binding of the F-Oda monomer to the PE with similar affinity as the BE even if different parts of the trimer are kept rigid.

To examine other complexes which appeared with significant probability, we summarized the structural ensemble using a rigid body transformation to describe the translation magnitude and rotation angle between the Oda et al. reference structure and the simulation structures (Fig. 2A). With this approach, we present the angle but do not specify the axis of rotation. This revealed that there were a total of six high probability regions (≥ 1%) (Fig. 2B), corresponding to six total high probability complexes (Fig. 2C). Probabilities were calculated as described in the Methods section. Movie S1 from a serial MD simulation shows transitions among some of these states. We found that there were two distinct structures at the BE, which corresponded to the first two peaks in the dRMS distributions (Fig. 1B). The higher probability complex is the complex which had lower dRMS, which is the more native- (filament) like of the two complexes (here called BE1). The other complex was significantly rotated with respect to the native-like filament BE complex by a rotation angle of 45-50 degrees (here called BE2). This complex involves contacts of the incoming monomer’s D-loop and subdomain 4 with the hydrophobic plug and subdomains 1, 3 of the two BE subunits of the dimer, respectively.

**Figure 2:**
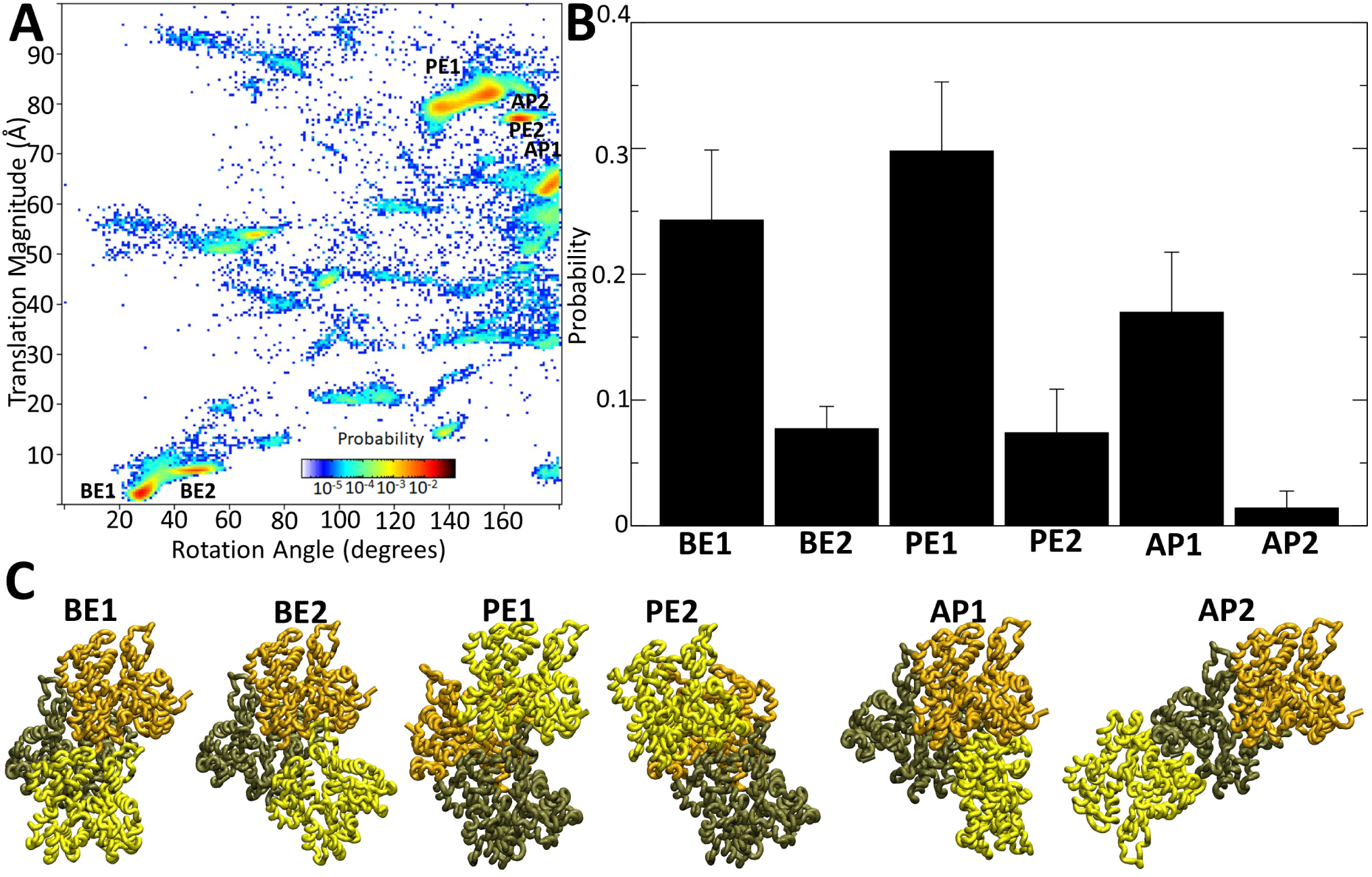
Simulation predicted structures of rigid F-Oda dimer plus monomer complex. (A) Plot of probability of finding the monomer at the corresponding translation and rotation configuration with respect to the BE bound state in Oda et al. (43) model. Data are for same simulations as Fig. 1B. (B) Probability of each high probability (>1%) state indicated in panel A, average and standard deviation from four separate simulations. (C) Images of structures in simulation hotspots shown in A.

The simulations gave two distinct complexes near the PE (Fig. 2). Again, the complex with higher probability (PE1) is the complex which was more filament-like (dRMS <5Å). The other complex (PE2) had high dRMS (appx. 20Å), suggesting it is in a notably different configuration than in the filament configuration. This complex could roughly be described by taking the long-pitch helix structure and shearing one subunit with respect to the other such that the subunit at the PE is shifted toward subdomain 4 of the other subunit (Fig. 2C). In this configuration, the D-loop of subunit *b* of the dimer is completely out of the hydrophobic groove of the monomer.

The complex AP1 (“antiparallel 1”), which formed in the simulations, bound at the BE at a very different configuration compared to BE1 or BE2 (Fig. 2). Instead of subdomains 2 and 4 making contact with the barbed end of subunit *c* of Fig. 1A, the incoming subunit *a* here could be described as “upside-down”, with subdomains 1 and 3 being the ones to make contacts, including with subunit *b*. This complex between the terminal BE subunit of the dimer and the incoming monomer bears some similarity to the antiparallel dimer, which has been proposed to have involvement in nucleation and polymerization (5, 16, 18) (Movie S2). This complex was not exactly the same as in crystal structure 1RFQ (16) (dRMS ∼ 14 Å); however, it is expected that the native antiparallel dimer should be notably different than the ones crystallized by cross-linking methods that further exhibited rotational variations dependent on crystal packing (16, 17). Complex AP2 is also an antiparallel complex but with the incoming subunit in antiparallel arrangement with respect to dimer subunit *b* rather than *c*.

### Contacts formed between actin subunits

To examine more precisely which residues were involved in stabilizing the various complexes we identified, we calculated the intermolecular contact map between the dimer and the monomer over the full simulation ensemble (Fig. 3A, B). To capture all regions which play a role in the formation of complexes, we used as a cutoff the value 2*σ* of the corresponding LJ interaction, to determine whether two residues were in contact (to capture even long range contacts that could be important in the formation of these complexes (57)). The residues with high contact probability are located at the pointed and barbed ends as well as the “back” side of the actin monomer, corresponding to the binding interfaces along the actin filament (Fig. 3B).

**Figure 3:**
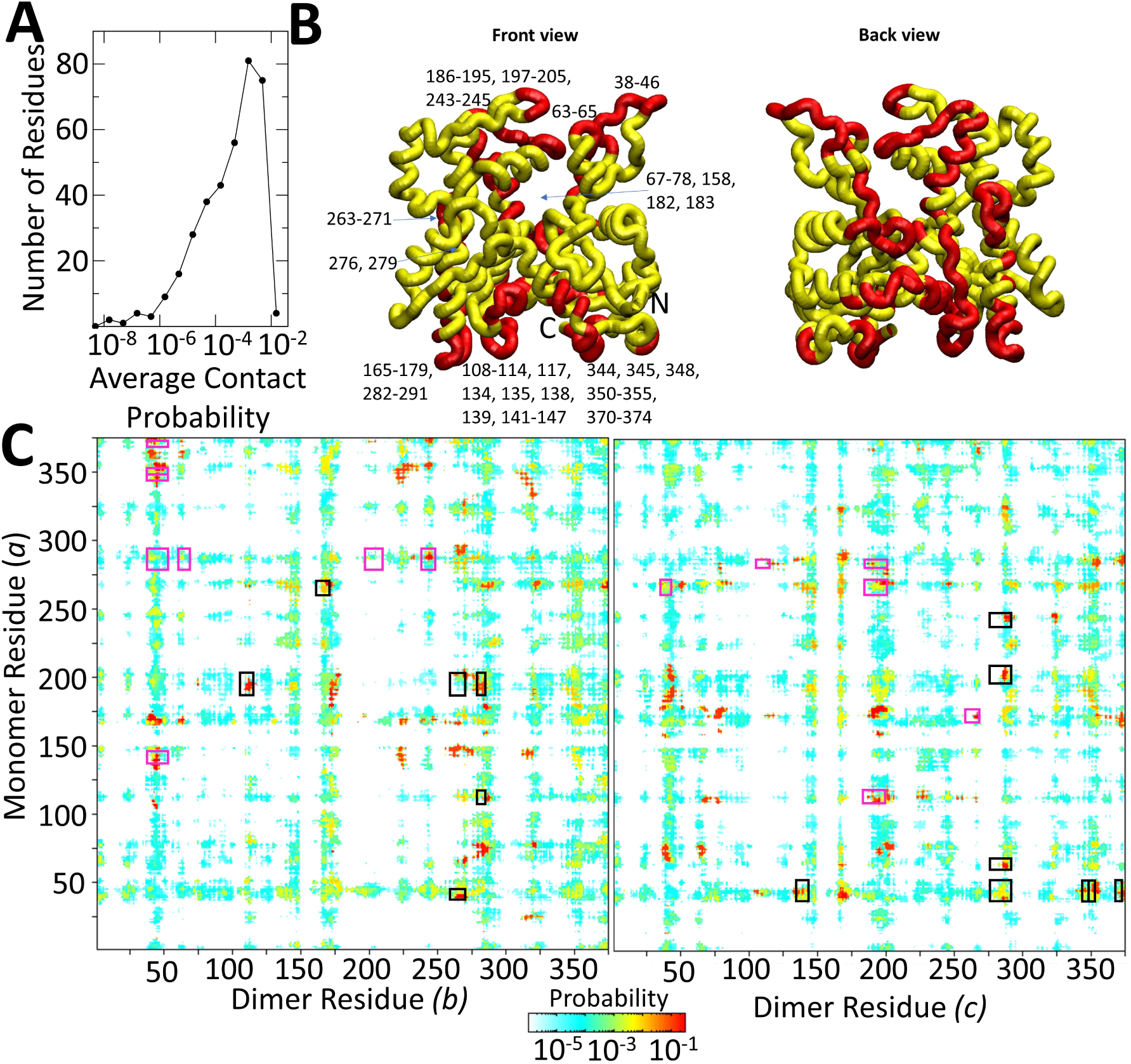
Contact probabilities of rigid F-Oda monomer with rigid F-Oda dimer. (A) Distribution of average contact probability over incoming monomer (subunit *a* in Fig. 1A) residues, for the same ensemble as Fig. 2. (B) Image highlighting in red the residues in the monomer with probability greater than 10^−3^ in the plot of panel A. (C) Contact map between incoming monomer and subunits *b* and *c* of dimer in Fig. 1A. Contact defined when residues are within 2*σ* of corresponding LJ potential of each other. Rectangles correspond to residues predicted to make contacts in the structure of Oda et al. (43). Black (pink) rectangles correspond to subunit bound at BE (PE)

Next we identified which residues in the incoming subunit are most important in interactions with the dimer. We calculated the average probability of contact of a given residue in the incoming monomer (subunit *a* in Fig. 1A) with all residues of each member of the dimer (subunits *b* and *c* in Fig. 1A), in the reference 197 K simulation (Fig. 3C). We found high probability regions of contact in most of the contact regions in the model of Oda et al. (43), shown by squares in Fig. 3C, see discussion in Supplemental Text.

### Simulation of actin polymerization mutants

To further evaluate the model, we performed simulations of actin mutated at surface resides that severely impair actin polymerization in vitro, without any significant structural changes of the C_*α*_ actin monomer backbone as studied by crystallography. These are the combined mutations K291E/P322K in subdomain 3 at the BE (58) and A204E/P243K in subdomain 4 at the PE (59–61). Each of these pair of mutations is located at the long-pitch helix contact interface.

The corresponding residue substitutions were introduced at the BE and PE of the rigid dimer, in both subunits *b* and *c* of Fig. 1A. The simulations of Fig. 1 and 2 were then repeated with an unmutated incoming monomer *a* interacting with the mutated dimer (we did not add the mutations to the incoming monomer to avoid clash of BE/PE mutations). We checked for the probabilities of BE1 (perturbed by K291E/P322K at the BE of the dimer) and PE1 (perturbed by A204E/P243K at the PE of of the dimer). The same six bound structures as Fig. 2 were observed. The dissociation constant of BE1 (influenced by K291E/P322K) increased by about 4-fold. This increase is consistent with a decreased ability for polymerization but we were not able to find experimental bounds on the critical concentration of the K291E/P322K mutant to compare quantitatively. However, the dissociation constant of state PE1 (influenced by A204E/P243K) did not not change appreciably compared to the unmutated case, even though the A204E/P243K mutation increases the critical concentration of polymerization from 0.1 to over 19 *µM* (59).

The relatively weak effect of the above mutations in the simulations indicates that the energies of the long-pitch contacts involving the corresponding aminoacids have been underestimated by the coarse-grained model in Fig. 1 and 2. Such single residue-level effects may be at the resolution limits of the KH model (as expected (31)) and/or require accounting for flexibility at the binding interface. Thus, Fig. 2B may underestimate the relative probability of BE1 and PE1, which, as native structures, may be strongly stabilized by specific contacts compared to the other states (for example, the side chain of M44 into the pocket above Y169 (9)).

### Effect of flexibility and conformation of incoming subunit

To gauge the importance of flexibility of the unstructured residues in actin, we repeated the same simulations of Figs. 1-3, but with flexibility in those residues (“F-Oda flex” model). We constrain parts of the dimer and monomer to be rigid as in previous simulations, but we now allow specific residues to be flexible, represented as beads connected along a chain by harmonic springs. We considered flexible the residues in the D-loop and near the N and C termini, which were missing in the crystal structure of G-ATP-actin (41). We did not add structural restraints such as angular or dihedral constraints and the D-loops at the pointed end of the dimer were kept rigid for reasons of computational efficiency (Fig. 4A).

**Figure 4:**
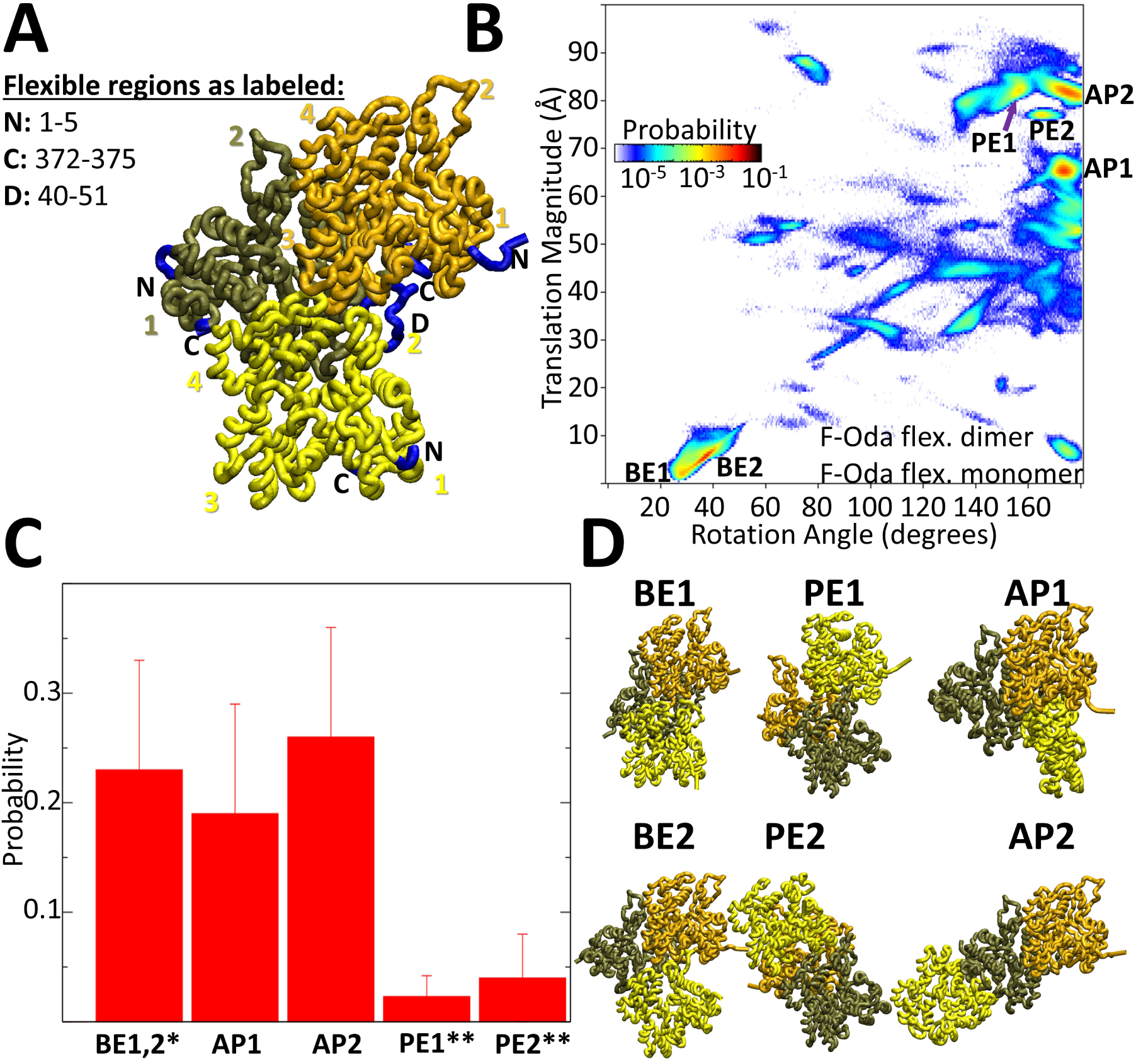
Effect of adding D-loop and N/C termini flexibility on structural ensemble of monomer interacting with dimer. (A) Cartoon showing the incoming monomer (subunit *a* in yellow) bound to the barbed end of dimer with subunits *b* and *c* connected to each other. Subunit *a* is in the F-Oda flex. configuration with residues made flexible at the termini and in the D-loop shown in blue. Subunits *b* and *c* have their termini flexible (blue) but the D-loop was kept rigid in the F-Oda configuration (Fig. 1). Other simulation parameters same as Fig. 1. (B) Rotation and translation distribution shown as in Fig. 2A. (C) High probability regions (>1%). Values show average and standard deviation from four separate simulations. *: BE1 and BE2 grouped together as they partially overlapped. **: Binding strength may be influenced by having only part of the PE binding interface flexible. (D) Images of high probability structures of panels B and C. The BE1 and BE2 images show structures at the two peaks of the common BE1/BE2 valley of attraction, connected by a continuum of high probability states.

The incoming F-Oda flex. monomer bound to the BE with high probability at configurations with rotation angles and translation magnitudes similar to the BE states in the F-Oda rigid case, so they were named the same, BE1 and BE2 (Fig. 4B-D). These BE1 and BE2 states were not however clearly separated now: monomers bound to the barbed spanned a range of angles between BE1 and BE2. Thus, the flexibility of the D-loop and C-termini allowed the incoming subunit to add to the barbed end over a range of angles. The modified contact probability map is shown in Fig. S3. In the simulations the D-loop fluctuated over a ranged of configurations, which were slightly more extended in an Oda-like conformation in the BE1 state (Fig. S4). The difference in dissociation constant of the F-Oda flex. monomer to the BE1 and BE2 states, with respect to the F-Oda rigid case, was not statistically significant.

The F-Oda flex. subunit also bound to the PE in PE1 and PE2 configurations, however with lower probability than the BE, a result that is likely influenced by keping the D-loops at the pointed end of the dimer rigid. Additionally, the F-Oda flex. simulations also indicated two regions which corresponded to antiparallel dimer-like configurations AP1 (Movie S2) and AP2. The combined AP1 and AP2 probability was greater than the BE probability, a result of the added flexibility when comparing to the rigid case where the reverse was observed. This flexibility influences both BE states and AP states that involve contacts between the flexible C termini.

We then examined the case of a monomer in the twisted G-actin conformation associating with an F-actin dimer, a situation that should more closely approximate the interaction of a filament with a monomer in the bulk. So we replaced the free monomer using the crystal structure of ATP G-actin of Graceffa et al. (41) (“G-ATP-Grac.”). The missing residues in the D-loop region and the N and C termini were added with the MODELLER software (42). These were the same residues that were made flexible in the F-Oda flex. case and were treated as flexible in the G-ATP-Grac. monomer as well. (Simulations were also conducted on a rigid G-actin with the missing residues added as shown in Fig. S5, to highlight the importance of the flexibility of these residues). The subunits of the dimer had flexible N and C termini and fixed D-loops as in Fig. 4A. Examples of serial MD simulations are shown in Movies S3-S5. The contact probability map is shown in Fig. S3.

We found that the G-ATP-Grac. monomer established three nearly-overlapping states of bound complexes at the BE (Fig. 5). Two of them were roughly the same BE complexes as in Figs. 2 and 4 (BE1 and BE2). For convenience we used the same BE1 and BE2 labels, even though the twisted configuration of G-actin does not allow the same long-pitch helix contacts in state BE1 as in Figs. 2 and 4. The third BE complex (BE3) also involved contacts with the D-loop of the incoming monomer with the dimer, with the monomer tilting out away from subunit *b* of the dimer. The BE3 complex had the highest probability of the BE complexes in this structural ensemble. In all BE1-BE3 states, the D-loop of incoming monomer *a* contacted subunit *c* of the dimer and showed similar hot spots in the contact map as in the F-Oda flex. monomer case (Fig. S3).

**Figure 5:**
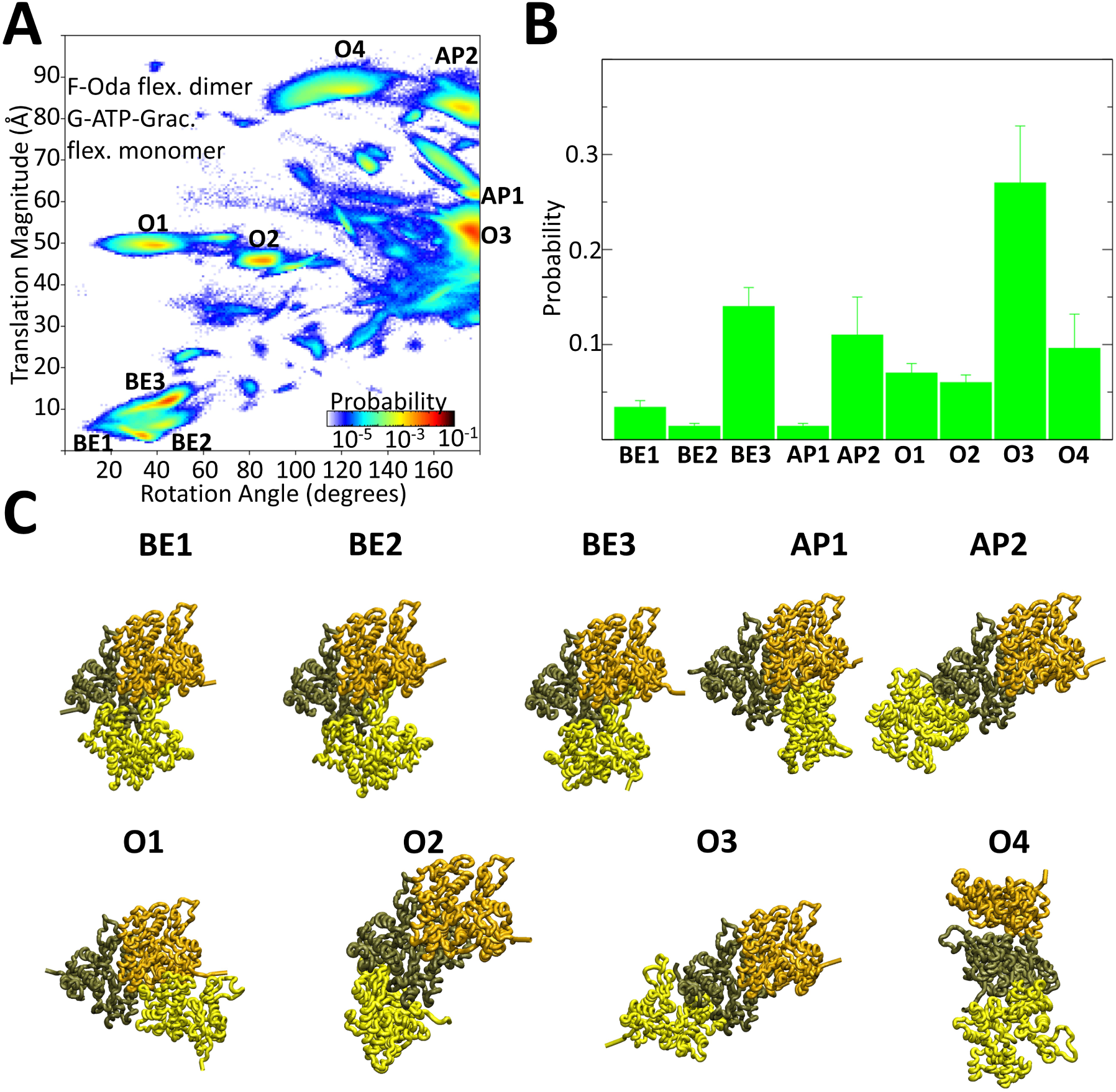
Interaction between G-actin monomer (G-ATP-Grac. flex) with a dimer in the F-Oda configuration. The same regions as in Fig. 4A were kept flexible in the monomer and dimer. Other simulation paramaters same as Fig. 1. (A) Rotation and translation distribution. (B) Probabilities of high probability regions (>1%). Values show average and standard deviation from four separate simulations. (C) Images of high probability structures of panels A and B.

The G-ATP-Grac. monomer formed AP1 (Fig. 5, Movie S2) and AP2 complexes (Fig. 5), similar to the F-Oda flex. monomer (Fig. 4), with AP2 now having higher probability. Interestingly however, the G-ATP-Grac. monomer did not bind at the pointed end. This observation provides support for the need of a conformational change of an incoming G-actin subunit for PE binding (8, 9), see Dicsussion.

Four additional complexes were observed with the G-ATP-Grac. monomer, named O1-O4 (O for “other” complex, Fig. 5). All four are structures near the BE, but in a configuration very different from the filament. In the O1 complex the back side of the incoming monomer contacted the front side of subunit *c* of the dimer. In structures O2, O3 and O4, the incoming monomer made contact with subunit *b* of the dimer. In O2 the incoming subunit is rotated nearly perpendicular with respect to the filament axis, and subdomains 2 and 4 of the incoming subunit are attached at the barbed end of subunit *b*. Complex O3 had probability much higher than any other complex. Complex O4 resembled an antiparallel structure, but the incoming subunit is rotated perpendicularly with respect to the axis of the filament. Complexes O1 and O3 bear some resemblance to a long-pitch helix configuration of incoming subunit *a* with either subunit *c* or *b*; however, instead of the two subunits being stacked on top of the other, there is a relative shift such that the that the faces of the subunits partially overlap.

### Interactions between two monomers

Having a basis for contact formation of a single actin monomer with a minimal filament, we then proceeded to study interactions between two monomers, which is important in understanding the mechanism of spontaneous filament nucleation. To accomplish this, we ran simulations with two actin monomers for the same three models for actin: F-Oda rigid, F-Oda flex., and G-ATP-Grac., the latter with flexible D-loop and termini (simulations were also conducted for rigid G-ATP-Grac. monomers; results presented in Fig. S5D). Specifically, we were interested in whether monomers would form short-pitch helix, long-pitch helix, and antiparallel dimer complexes. The total probability of each complex was measured by calculating the dRMS with respect to each complex and measuring the magnitude of the smallest peak in the dRMS distribution (Fig. 6). For the F-Oda rigid monomers, each of the three complexes were identified as a peak at small dRMS; however, the probability of the long-pitch helix complex was low (<1%). In the flexible cases, we did not detect any clear long-pitch helix complex formation; the smallest dRMS complex corresponded to the complexes shown in Fig. 6 and had low probability (≪ 1%). Both the short-pitch helix and the antiparallel complexes were detected with high probability. The flexibility in the D-loop led to its wrapping around the side of the other subunit in the short-pitch dimer.

**Figure 6:**
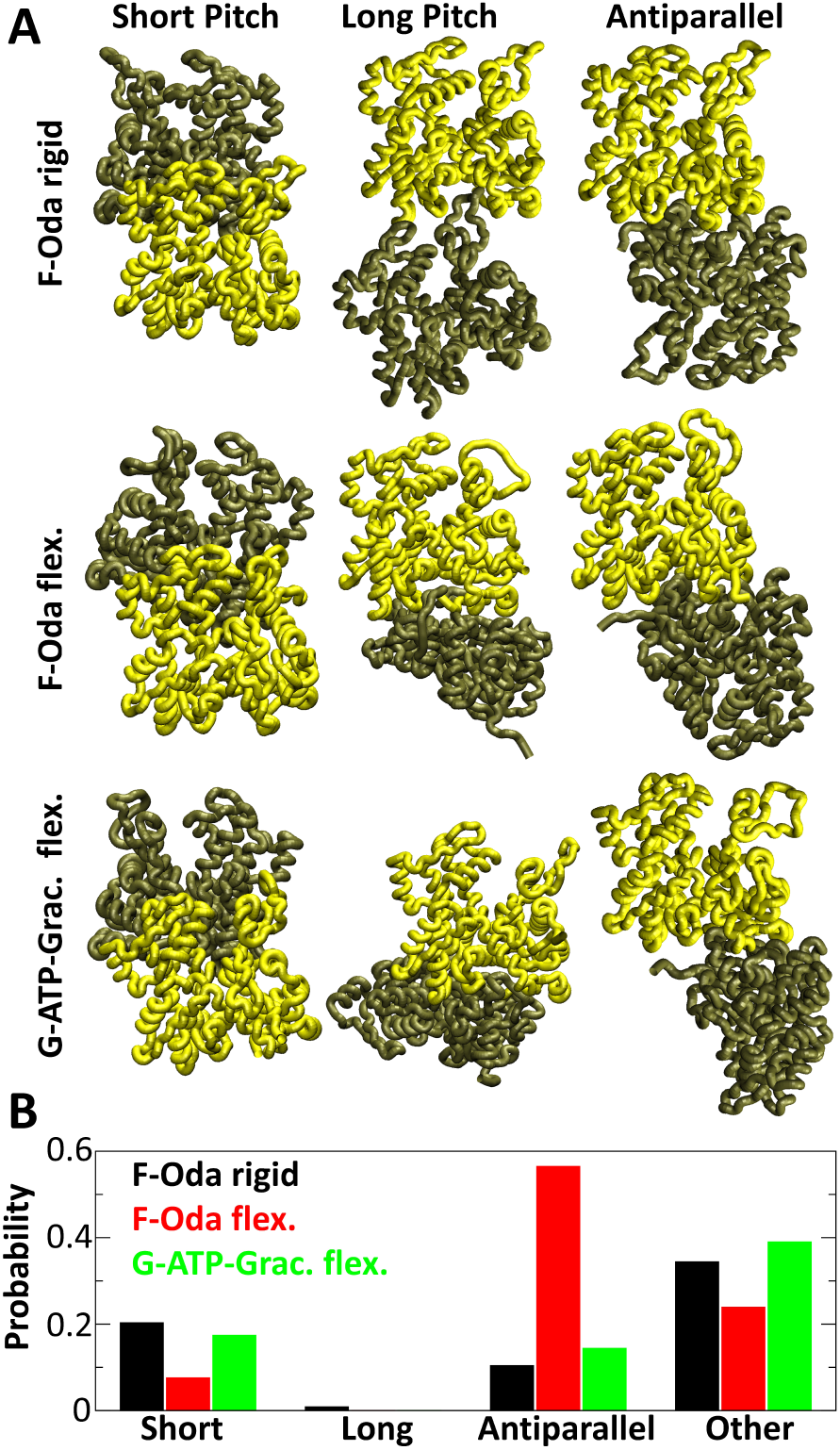
Structures and probabilities in monomer-monomer binding. Results of simulations with two monomers in the rigid F-Oda, F-Oda flex., and G-ATP-Grac. flex. configurations that correspond to subunit *a* in Fig. 1, 4 and 5, respectively. Other simulation parameters same as Fig. 1, except box size is 300 Å in each direction for simulations with F-Oda monomers and 149 Å for G-ATP-Grac. flex. monomers. (A) Images of the lowest peak in the dRMS distribution with respect to the respective reference structure (short pitch, long pitch, and antiparallel). For the short-pitch helix structures, the dRMS peak occurred at less than 5 Å. For the long-pitch helix, the peak was at 5 Å for F-Oda rigid and at 8 Å for the other two cases. For the antiparallel case, dRMS peaks in vertical ordering were at 12, 7, and 5 Å. (B) Probability comparison of the structures shown in A. Other structures shown in Fig. S7. F-Oda actin monomer simulations performed at concentration of 1*µ*M and G-ATP-Grac monomer simulations performed at concentration of 1mM.

Both the flexibility in the termini and the conformation of the subunit (G-actin as opposed to F-actin) contributed to lowering of the dRMS of the antiparallel dimer complex with respect to the 1RFQ crystal structure (16). This is realized by a rotation of subdomains 2 and 4 of the bottom subunit of Fig. 6 into the page (see Fig. S6).

As with the previous simulations, other high probability complexes formed in addition to the short-pitch helix, long-pitch helix, and antiparallel dimer (Fig. S7). Some of these complexes involve contacts with regions of the monomer that were previously buried insider the dimer (structures F1, F2, F5, T1, T2, T3 in Fig. S7). The F1 and F2 structures were the main new complexes between F-Oda and F-Oda flex. that laid with their flattened sides against each other. The main other complex between G-ATP-Grac. monomers (probability 31.4%) resembled the O1 and O3 structures of Fig. 5 (where O1 and O3 are similar structures that the incoming monomer makes separately with each subunit of the dimer). The O1/O3 structure resembles a displaced long-pitch complex and thus it may play a role in nucleation. Most of the other structures in Fig. S7 differed significantly from an F-actin arrangement, except for T1 that has a dRMS ≈ 13 Å with respect to a short-pitch dimer.

The low probability or absence of long-pitch helix contacts in comparison to the short-pitch helix in Fig. 6 and S6 is in contradiction with the expectation that these are the dominant contacts during nucleation and along the actin filament (5, 9, 19, 28, 49), a point discussed further in the Discussion section.

### Interactions between many subunits: filament stability and polymerization

Given the model’s success in predicting association of a monomer to the BE or PE, we examined the potential of the model to simulate polymerization of multiple subunits. First we asked whether the model would hold an actin filament stable over the course of a simulation at fixed temperature. Starting with a filament of 6 rigid F-Oda subunits arranged in the Oda et al. configuration, we found that there were minor rearrangements over the course of a simulation, but no subunits fell off of the filament (Fig. 7A, Fig. S8A). Then we ran serial simulations beginning from a pool of rigid F-Oda monomers to see if they would come together to form a filament on timescales easily accessible by simulation. A slightly larger temperature of 204 K compared to 197 K (corresponding to a modest increase in the *K_d_* of bound states by factor 3.4) was used to allow monomers to reconfigure more easily out of metastable minima that occur in serial simulation while still maintaining the same binding states. We found that the monomers did form a filament nucleus that polymerized into a short filament with most subunits in the standard filament configurations (Fig. 7B, S8B). Subunits were able to fall off of the PE of the filament, however a clump of actin subunits typically localized to the BE of the filament. In the Discussion section we briefly discuss if such aggregates and the AP1 or AP2 structures may occur during actin polymerization.

**Figure 7:**
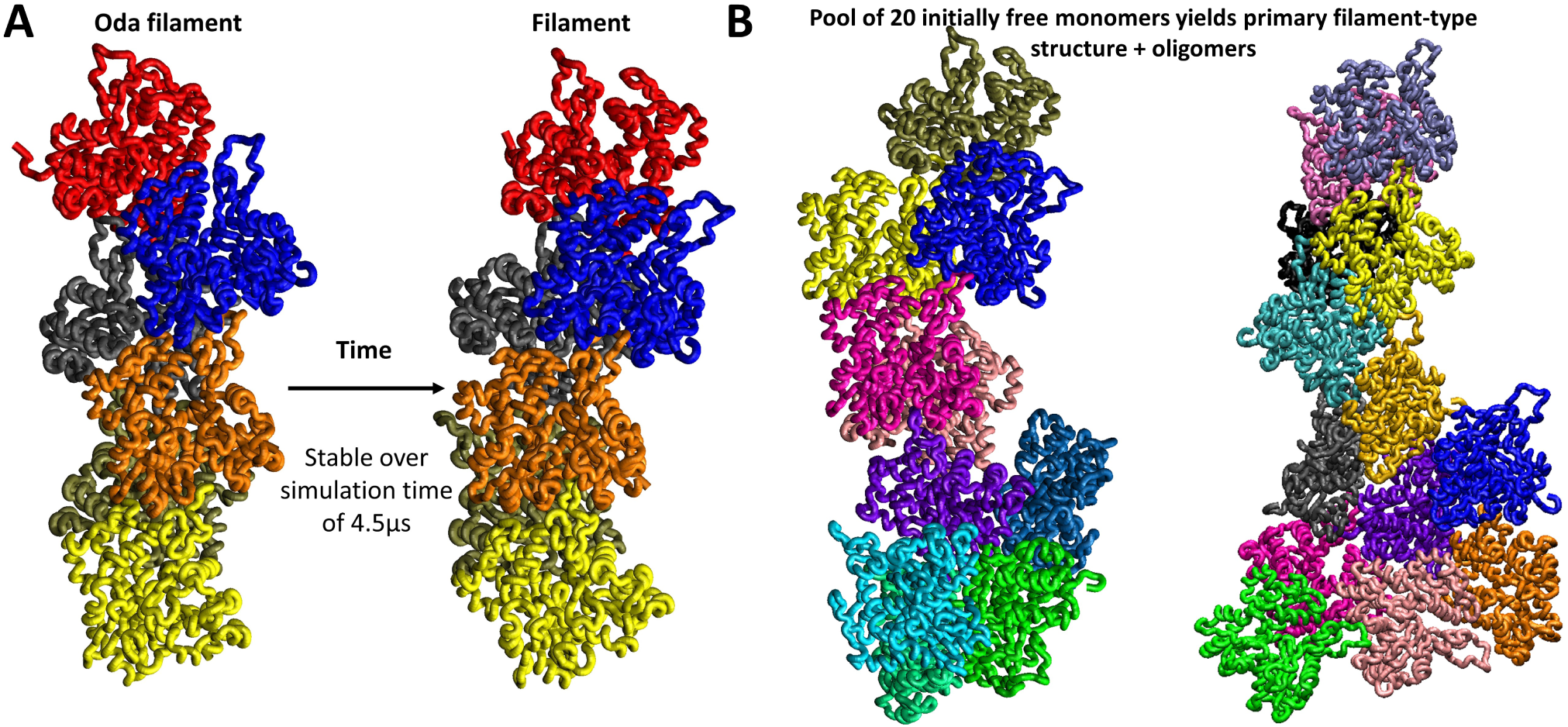
Simulations of filament stability and polymerization. (A) Serial MD simulation (Model A with 3*σ* cutoff at 197 K) starting from 6 rigid F-Oda monomers placed in the Oda et al. (43) filament configuration. The filament remains stable over the course of a serial simulation. (B) Serial MD simulation of a pool of rigid F-Oda monomers dispersed in a box of size 1500 Å^3^ (Model A with 3*σ* cutoff at 204 K, concentration 9.84 *µM*) yields primary filament-type structures with aggregates forming at the the barbed end. Examples of two simulations after at least 1 *µs* (in reduced simulation time that does not map to physical time).

To further examine the pathway of addition of a new subunit, we examined the ability of a rigid F-Oda subunit at the BE of a rigid F-Oda dimer to transition between the different complexes by running standard serial MD simulations (Movie S1). We found that it was common for the subunit to transition between BE2 and BE1 multiple times before dissociation (Movie S1); after dissociation, the subunit could again later bind the BE (Fig. S9). The incoming subunit reached the polymerizable BE1 state either directly or through the intermediate BE2 state (Fig. S9). We also performed standard serial MD simulations with the G-ATP-Grac. flex. monomer binding to the F-Oda flex. dimer and also found it common for there to be multiple transitions between the BE states before dissociation (Movie S3). Additionally we observed association events directly to the BE. We found the D-loop may initially bind the dimer and help the incoming subunit fall into a BE state (Movie S4) or both subdomains 2 and 4 of the incoming subunit will stick to the dimer and eventually slide into a BE configuration (Movie S5).

## 4 DISCUSSION

In this work, we applied a coarse-grained model of multiprotein complex formation to actin polymerization. We used the KH model, which was previously validated on several protein complexes (31). The KH model successfully predicted transient protein encounter complexes (32), though refinement through NMR experimental measurements was necessary to reproduce the correct population of non-specific complexes. Since we did not perform experimental measurements as part of this investigation, and no data exist to precisely distinguish among encounter complexes, our results should be interpreted as an initial step towards a future, more accurate, coarse-grained model for actin polymerization and nucleation. Such a coarse-grained model would need further testing and refinement against experimental data and to further consider conformational transitions such as propeller twist and cleft opening/closing (5, 6) as well as the structural polymorphism of the actin filament (27, 47). Comparison to other coarse-graining approaches that can predict protein association is also an area for future investigation (62, 63).

The model was successful in reproducing binding of an actin monomer at the anticipated location at barbed and pointed ends of a short actin filament, without any parameter tuning. Interactions along the short-pitch direction were stronger than the long pitch, between all monomer configurations that we explored (Fig. 6). This is largely due to the fact that in this model, the number of pairs of aminoacids participating in short-pitch contacts (as defined in Fig. 3) are more numerous that those along the long-pitch direction (313 ± 27 vs. 119 ± 25 between an incoming monomer and filament). By contrast, measurement of contact surface area (9, 49), energy calculations (28), and lateral intrastrand slipping in electron micrographs (52), have instead suggested that the long-pitch helix interaction is stronger compared to the short pitch. Future work, including all atom simulations, should help clarify the energetics of the two types of contacts.

One aspect of the coarse-grained model that may require adjustment is the enhancement of the binding strength along the long-pitch helix direction. One indication of this is the comparatively weak effect of polymerization mutations K291E/P322K and A204E/P243K between subdomains 3 and 4 in the model. The complex electrostatic interactions of the highly charged actin molecule may also contribute to this effect (40). It has been reported that the actin polymerization Δ*H* changes sign and Δ*S* decreases 60-fold as the KCl concentration increases from 30 to 100 mM (56), at which point the critical concentration exhibits a minimum (30). An important factor that we did not include, and which may play a role in enhancing long-pitch helix contacts, is the divalent cation binding at the “polymerization cation site” between subdomains 3 and 4 or at the “stiffness cation site” within the D-loop interface (40). Enhancing long-pitch contacts would also enhance overall binding affinity to the barbed and pointed ends, bringing the coarse-grained model’s *K_d_* for the barbed end closer to the measured value of the critical concentration.

The model reproduced binding of actin subunits in the antiparallel configuration, as observed prior experiments (14–16, 18, 19, 64). Intriguingly, the model predicts that the AP configuration can also occur at the barbed end (states AP1/AP2). Providing an accurate estimate of the AP1 *K_d_* is difficult, given the above stated uncertainties in calibrating for the correct long-pitch helix contact energetics. Kinetic studies of actin nucleation using SAXS suggested formation of non-polymerizable dimers with an equilibrium dissociation constant 160 *µM* (19). If we use this value as a reference for the dissociation constant of the AP2 state or the AP state between two monomers, the estimated value of the AP1 *K_d_* varies significantly depending on the model (Figs. 2, 4, 5 and Table S1) between 2 to 1000 *µM*. The rate of actin polymerization is linear as function of actin monomer concentration, up to 20 *µM* (7), suggesting that either the AP1 barbed end state is very short lived (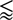1 ms) or else the *K_d_* for the AP1 state is higher than 20 *µM*. Alternatively, monomers in the AP1 state might be able to transition to a polymerizable state without leaving the barbed end. Indeed, in serial simulations we frequently observed transitions from AP1 to BE1/BE2 states through reorientation of a monomer weakly bound to the barbed end (Movies S1, S3). The AP state might be important under cellular conditions where unpolymerized actin exists at ~ 100 *µM* (65). Profilin, which binds at the barbed end of a big fraction of unpolymerized actin would prevent AP state formation and thus may have another role to play in regulating polymerization.

Even though our study focused on equilibrium properties, our simulations also provide insights into the kinetics of actin polymerization. Elongation requires about 50 collisions of an incoming actin monomer with the barbed end (10). The dependence of actin polymerization rate on viscosity implies diffusion-controlled kinetics (10), which has been thought to suggest limitation by actin monomer diffusing to the correct orientation (66), with the possible aid of electrostatic steering (29). In our simulations, the incoming monomer bound to the barbed end over a range of angles that involved pivoting around the D-loop making contacts with penultimate subunit at the barbed end. This orientational variability was a robust feature of all three models used to describe polymerization (Fig. 2,4 and 5). We suggest that transition between these states reflects a process of searching for the correct orientation after diffusion near the end (67). Other plausible polymerization pathways might involve the O1 state of Fig. 5 that resembles a displaced long-pitch helix contact (Movie S5).

Further information on the mechanism of polymerization is provided by the observation of transitions between encounter complexes at either end in serial MD simulations (Movies S1, S3-S5, which were however performed at reduced viscosity to enable efficient sampling of bound states). The kinetics in these movies suggest that the addition of a new G-actin subunit to the barbed end could indeed involve initial contact of the D-loop of the incoming monomer, followed by pivoting around the contact point along angles spanning the BE1/BE2/BE3 states. Flattening of this incoming subunit into an F-actin form, an effect that is not included in our current simulations, would allow better long-pitch helix contacts of both the D-loop and subdomain 3 of the incoming monomers (as in the BE1 state of the F-Oda incoming monomers in Figs. 2 and 4). In this pathway, the longitudinal contact between subdomain 3 of the incoming monomer with subdomain 4 of the filament subunit along the long-pitch helix direction is established in a second step. This is the reverse order of a pathway suggested based on structural studies (9, 49). In our proposed pathway, the flexibility of the D-loop would increase the capture radius of the diffusive search process and enhance the polymerization rate constant. Such a polymerization mechanism could explain why ADP-actin, whose D-loop has been proposed (though not conclusively) to have a higher tendency to fold into an alpha helix (6, 27, 68, 69), has a polymerization rate constant that is 3-fold lower compared to ATP-actin (70).

Irrespective of the pathway of monomer arrival to the barbed end, our study further clarifies how flattening of the actin subunit contributes to polymerization and filament stability (5, 6, 9). In states BE1, BE2, and BE3 of Fig. 6, the incoming G-actin subunit adopts a tilted conformation that prevents it from making the same long-pitch helix contacts as the F-actin subunit in state BE1 of Fig. 2. Flattening of the subunit would enable these additional contacts and enhance filament stability (53).

Our study motivates future work to test if the barbed end polymerization search process is influenced by transient encounter complexes (71) and/or transient barbed end aggregates (Fig. 7) in both productive and unproductive ways. Proteins that associate to the barbed during polymerization, such as profilin, formins and Ena/VASP might not only regulate polymerization through enhancement of local concentration and orientation steering but also by preventing unproductive complexes that could become limiting at cellular elongation rates. Methods such as paramagnetic relaxation enhancement nuclear magnetic resonance spectroscopy in combination with modeling (32, 71), could clarify these issues. Further analysis is also needed to determine the relevance of such polymerization/depolymerization pathways to length fluctuations at steady state in vitro (12, 13, 20).

The lack of binding of G-actin to the pointed end in the simulations of Fig. 5 (despite binding of a monomer in the F-actin conformation in Figs. 2 and 4) provides further support for polymerization at pointed end being partly limited by conformational change of the incoming monomer (8, 9). This limit is in addition to conformational changes required by the the terminal subunits at the pointed end (8, 9), which we did not explore: in this study we assumed that the terminal barbed and pointed end subunits have the same structure as the filament interior; this assumption is expected to be a good approximation for the barbed but not the pointed end (8, 9).

The difference in the critical concentration between the two ends of the actin filament is a fundamental property of actin, underlying the treadmilling of single filaments in vitro (20). In a model proposed to explain the difference in the critical concentration, which also takes into account microscopic reversibility, it was important that the rate constant for addition of ATP-actin to an ADP-actin terminal subunit is lower than that to an ATP-actin terminal subunit (70). We speculate that this difference may be related to conformational differences, including D-loop flexibility, between ADP-actin and ATP-actin at the terminal subunits of the pointed end (ADP-Pi-actin being short lived (70)).

Finally, we discuss the implications of our study on the mechanism of actin filament nucleation. In our simulations, monomers in the F-Oda rigid or G-ATP-Grac. flex. configuration established filamentous contacts preferentially along the short-pitch helix direction (Fig. 6). This binding between monomers occurred with a *K_d_* that was approximately 400 times higher than the combined *K_d_* of the same type of monomer to the barbed end states BE1, BE2, and BE3. The large value of the *K_d_* ratio is in agreement with theories of actin filament nucleation in which dimers are highly unstable intermediates to a stable filament nucleus that is a trimer or tetramer (19, 28). Quantitatively, this *K_d_* ratio is near the lower range of prior models of actin filament nucleation, summarized in (19). Our estimate of *K_d_* enhancement is closer to that in recent modeling and SAXS experiments by Oda et al. (19) (500 times higher) than in the model of Sept and McCammon (28) (3.5 10^5^ times higher), noting that in both cases the long-pitch helix dimer was the one expected to be more stable than the short-pitch helix dimer (even though G-actin cannot make the same type of long-pitch helix contacts as F-actin due to its twist).

Our work further supports the presence of non-filamentous contacts between actin monomers, in agreement with modeling of SAXS observations (19). We observed formation of antiparallel dimers (19) as well as multiple other complexes (Fig. S7). One notable complex is the O1/O3 configuration (Fig. 5 and S7) that occurred with high probability in our simulation with G-actin and resembled a displaced long-pitch helix dimer. It’s plausible that actin nucleation is limited by the shift of this O1/O3 state to a regular F-actin long-pitch helix with the help of additional short-pitch helix contacts of a trimer. Future kinetic studies that extend our coarse-grained model to include conformational changes of the actin monomers could test the proposed activation step during nucleation (19), the role of antiparallel dimer in nucleation (15), and other transient encounter complexes such as O1/O3.

In summary, coarse-grained modeling provides insights into intermediates formed during actin nucleation and polymerization in a way that would currently be difficult to achieve using all-atom simulations. Building such models is a critical first step in building a deeper understanding of molecular mechanisms behind actin polymerization and its regulation by other proteins, such as profilin, Arp2/3 complex, formins and other nucleation and elongation promoting factors.

## Supporting information

LAMMPS input

## AUTHOR CONTRIBUTIONS

BH and DV contributed to design of research, interpretation of results and wrote the paper. BH performed simulations and analysis with the input of DV and Jeetain Mittal (who declined to be co-author). AH performed analysis and validation simulations with input from BH and DV.

## ACKNOWLEDGMENTS

We thank Jeetain Mittal for his contributions to design of the research, design of figures, and interpretation of results. We also thank Young C. Kim for help with validation and for providing computational methods, and Thomas Pollard for detailed feedback on the manuscript. This work was supported by National Institutes of Health Grant R01GM114201. Use of the high-performance computing capabilities of the Extreme Science and Engineering Discovery Environment (XSEDE), which is supported by the National Science Foundation, project no. TG-MCB180021 is also gratefully acknowledged.

## SUPPORTING CITATIONS

References (72, 73) appear in the Supporting Material.

## REFERENCES

1. Laurent Blanchoin, Rajaa Boujemaa-Paterski, Cecile Sykes, and Julie Plastino. “Actin dynamics, architecture, and mechanics in cell motility.” eng. In: Physiol Rev 94.1 (2014), pp. 235–263.

2. Fumio Oosawa. “My various thoughts on actin”. In: *Biophysics and Physicobiology* 15 (2018), pp. 151–158.

3. Thomas D. Pollard. “Actin and Actin-Binding Proteins”. In: Cold Spring Harbor Perspectives in Biology 8.8 (2016).

4. Marie-France Carlier, Julien Pernier, Pierre Montaville, Shashank Shekhar, Sonja Kühn, and Cytoskeleton Dynamics and Motility group. “Control of polarized assembly of actin filaments in cell motility”. In: Cellular and Molecular Life Sciences 72.16 (2015), pp. 3051–3067.

5. Roberto Dominguez and Kenneth C. Holmes. “Actin Structure and Function”. In: Annual Review of Biophysics 40.1 (2011), pp. 169–186.

6. Dmitri S. Kudryashov and Emil Reisler. “ATP and ADP actin states”. In: Biopolymers 99.4 (2013), pp. 245–256.

7. Thomas D. Pollard. “Rate Constants for the Reactions of ATP- and ADP-Actin with the Ends of Actin Filaments”. In: Journal of Cell Biology 103 (1986), pp. 2747–2754.

8. Akihiro Narita, Toshiro Oda, and Yuichiro Maéda. “Structural basis for the slow dynamics of the actin filament pointed end”. In: The EMBO Journal 30.7 (2011), pp. 1230–1237.

9. Steven Z. Chou and Thomas D. Pollard. “Mechanism of actin polymerization revealed by cryo-EM structures of actin filaments with three different bound nucleotides”. In: Proceedings of the National Academy of Sciences 116.10 (2019), pp. 4265–4274.

10. D Drenckhahn and T D Pollard. “Elongation of actin filaments is a diffusion-limited reaction at the barbed end and is accelerated by inert macromolecules.” In: Journal of Biological Chemistry 261.27 (1986), pp. 12754–12758.

11. T. L. Hill. Linear aggregation theory in cell biology. Springer series in molecular biology. New York: Springer-Verlag, 1987.

12. D. Vavylonis, Q. Yang, and B. O’Shaughnessy. “Actin polymerization kinetics, cap structure, and fluctuations”. In: Proc. Natl. Acad. Sci. U. S. A. 102 (2005), pp. 8543–5488.

13. E. B. Stukalin and A. B. Kolomeisky. “ATP Hydrolysis Stimulates Large Length Fluctuations in Single Actin Filaments”. In: Biophys. J. 90 (2006), pp. 2673–2685.

14. R Millonig, H Salvo, and U Aebi. “Probing actin polymerization by intermolecular cross-linking.” In: The Journal of Cell Biology 106.3 (1988), pp. 785–796.

15. Michael R. Bubb, Lakshmanan Govindasamy, Elena G. Yarmola, Sergey M. Vorobiev, Steven C. Almo, Thayumanasamy Somasundaram, Michael S. Chapman, Mavis Agbandje-McKenna, and Robert McKenna. “Polylysine Induces an Antiparallel Actin Dimer That Nucleates Filament Assembly: Crystal Structure at 3.5-Å Resolution”. In: Journal of Biological Chemistry 277.23 (2002), pp. 20999–21006.

16. Robbie Reutzel, Craig Yoshioka, Lakshamanan Govindasamy, Elena G. Yarmola, Mavis Agbandje-McKenna, Michael R. Bubb, and Robert McKenna. “Actin crystal dynamics: structural implications for F-actin nucleation, polymerization, and branching mediated by the anti-parallel dimer”. In: Journal of Structural Biology 146 (2004), pp. 291–301.

17. Vadim A. Klenchin, Sofia Y. Khaitlina, and Ivan Rayment. “Crystal Structure of Polymerization-Competent Actin”. In: Journal of Molecular Biology 362.1 (2006), pp. 140–150.

18. Unai Silván, Céline Boiteux, Rosmarie Sütterlin, Ulrich Schroeder, Hans Georg Mannherz, Brigitte M. Jockush, Simon Bernèche, Ueli Aebi, and Cora-Ann Schoenenberger. “An antiparallel actin dimer is associated with the endocytic pathway in mammalian cells”. In: Journal of Structural Biology 177 (2012), pp. 70–80.

19. Toshiro Oda, Tomoki Aihara, and Katsuzo Wakabayashi. “Early nucleation events in the polymerization of actin, probed by time-resolved small-angle x-ray scattering”. In: Scientific Reports 6 (2016), p. 34539.

20. I. Fujiwara, S. Takahashi, H. Tadakuma, T. Funatsu, and S. Ishiwata. “Microscopic analysis of polymerization dynamics with individual actin filaments.” In: Nat. Cell Biol. 4 (2002), pp. 666–673.

21. Jhih-Wei Chu and Gregory A. Voth. “Coarse-Grained Modeling of the Actin Filament Derived from Atomistic-Scale Simulations”. In: Biophysical Journal 90 (2006), pp. 1572–1582.

22. Jim Pfaendtner, Davide Branduari, Michele Parrinello, Thomas D. Pollard, and Gregory A. Voth. “Nucleotide-dependent conformational states of actin”. In: PNAS 106 (2009), pp. 12723–12728.

23. Marissa G. Saunders and Gregory A. Voth. “Comparison between Actin Filament Models: Coarse-Graining Reveals Essential Differences”. In: Structure 20 (2012), pp. 641–653.

24. Marco A. Deriu, Ardita Shkurti, Giulia Paciello, Tamara C. Bidone, Umberto Morbiducci, Elisa Ficarra, Alberto Audenino, and Andrea Acquaviva. “Multiscale modeling of cellular actin filaments:From atomistic molecular to coarse-grained dynamics”. In: Proteins: Structure, Function & Bioinformatics 80 (2012), pp. 1598–1609.

25. Osman N. Yogurtcu, Jin Seob Kim, and Sean X. Sun. “A Mechanochemical Model of Actin Filaments”. In: Biophysical Journal 103.4 (2012), pp. 719–727.

26. Behzad Mehrafrooz and Amir Shamloo. “Mechanical differences between ATP and ADP actin states: A molecular dynamics study”. In: Journal of Theoretical Biology 448 (2018), pp. 94–103.

27. Fikret Aydin, Harshwardhan H. Katkar, and Gregory A. Voth. “Multiscale simulation of actin filaments and actin-associated proteins”. In: Biophysical Reviews 10 (2018), pp. 1521–1535.

28. David Sept and J. Andrew McCammon. “Thermodynamics and kinetics of actin filament nucleation”. In: Biophysical Journal 81 (2001), pp. 667–674.

29. David Sept, Adrian H Elcock, and J.Andrew McCammon. “Computer simulations of actin polymerization can explain the barbed-pointed end asymmetry”. In: Journal of Molecular Biology 294.5 (1999), pp. 1181–1189. issn: 0022-2836.

30. Jun Ohnuki, Akira Yodogawa, and Mitsunori Takano. “Electrostatic balance between global repulsion and local attraction in reentrant polymerization of actin”. In: Cytoskeleton 74 (2017), pp. 504–511.

31. Gerhard Hummer and Young Kim. “Coarse-grained Models for Simulations of Multiprotein Complexes: Application to Ubiquitin Binding”. In: Journal of Molecular Biology 375 (2008), pp. 1416–1433.

32. Young C. Kim, Chun Tang, G. Marius Clore, and Gerhard Hummer. “Replica exchange simulations of transient encounter complexes in protein–protein association”. In: Proceedings of the National Academy of Sciences 105.35 (2008), pp. 12855– 12860.

33. Brandon G. Horan, Gül H. Zerze, Young C. Kim, Dimitrios Vavylonis, and Jeetain Mittal. “Computational modeling highlights the role of the disordered Formin Homology 1 domain in profilin-actin transfer”. In: FEBS Lett. 592 (2018), pp. 1804–1816.

34. Sanzo Miyazawa and Robert L. Jernigan. “Residue – Residue Potentials with a Favorable Contact Pair Term and an Unfavorable High Packing Density Term, for Simulation and Threading”. In: Journal of Molecular Biology 256.3 (1996), pp. 623–644.

35. Steven J. Plimpton. “Fast Parallel Algorithms for Short-Range Molecular Dynamics”. In: FEBS Letters 592 (2018), pp. 1804–1816.

36. Pieter J. in ’t Veld, Steven J. Plimpton, and Gary S. Grest. “Accurate and efficient methods for modeling colloidal mixtures in an explicit solvent using molecular dynamics”. In: Computer Physics Communications 179 (2008), pp. 320–329.

37. Robert H. Swendsen and Jian-Sheng Wang. “Replica Monte Carlo Simulation of Spin-Glasses”. In: Physical Review Letters 57 (1986), pp. 2607–2609.

38. Enrique M. De La Cruz, Anna Mandinova, Michel O. Steinmetz, Daniel Stoffler, Ueli Aebi, and Thomas D. Pollard. “Polymerization and structure of nucleotide-free actin filaments11Edited by W. Baumeister”. In: Journal of Molecular Biology 295.3 (2000), pp. 517–526.

39. Hyeran Kang, Michael J. Bradley, Brannon R. McCullough, Anaëlle Pierre, Elena E. Grintsevich, Emil Reisler, and Enrique M. De La Cruz. “Identification of cation-binding sites on actin that drive polymerization and modulate bending stiffness”. In: Proceedings of the National Academy of Sciences 109.42 (2012), pp. 16923–16927.

40. Hyeran Kang, Michael J. Bradley, W. Austin Elam, and Enrique M. De La Cruz. “Regulation of Actin by Ion-Linked Equilibria”. In: Biophysical Journal 105.12 (2013), pp. 2621–2628. issn: 0006-3495.

41. Phillip Graceffa and Roberto Dominguez. “Crystal structure of monomeric actin in the ATP state. Structural basis of nucleotide-dependent actin dynamics”. In: Journal of Biological Chemistry 278 (2003), pp. 34172–34180.

42. Benjamin Webb and Andrej Sali. “Comparative Protein Structure Modeling Using MODELLER”. In: Current Protocols in Bioinformatics 54 (2016), pp. 1–37.

43. Toshiro Oda, Mitsusada Iwasa, Yuchiro Maéda, and Akihiro Narita. “The nature of globular- to fibrous-actin transition”. In: Nature 457 (2009), pp. 441–445.

44. Dimitrios Vavylonis, David R. Kovar, Ben O’Shaughessy, and Thomas D. Pollard. “Model of formin-associated actin filament elongation”. In: Molecular Cell 21 (2006), pp. 455–466.

45. Takashi Fujii, Atsuko H. Iwane, Toshio Yanagida, and Keiichi Namba. “Direct visualization of secondary structures of F-actin by electron cryomicroscopy”. In: Nature 467 (2010), p. 724.

46. Kenji Murakami, Takuo Yasunaga, Taro Q.P. Noguchi, Yuki Gomibuchi, Kien X. Ngo, Taro Q.P. Uyeda, and Takeyuki Wakabayashi. “Structural Basis for Actin Assembly, Activation of ATP Hydrolysis, and Delayed Phosphate Release”. In: Cell 143.2 (2010), pp. 275–287.

47. Vitold E. Galkin, Albina Orlova, Gunnar F. Schröder, and Edward H. Egelman. “Structural polymorphism in F-actin”. In: Nature Structural & Molecular Biology 17 (2010), p. 1318.

48. Julian von der Ecken, Mirco Müller, William Lehman, Dietmar J. Manstein, Pawel A. Penczek, and Stefan Raunser. “Structure of the F-actin-tropomyosin complex”. In: Nature 519 (2014), p. 114.

49. Vitold E. Galkin, Albina Orlova, Matthijn R. Vos, Gunnar F. Schröder, and Edward H. Egelman. “Near-Atomic Resolution for One State of F-Actin”. In: Structure 23.1 (2015), pp. 173–182. issn: 0969-2126.

50. Felipe Merino, Sabrina Pospich, Johanna Funk, Thorsten Wagner, Florian Küllmer, Hans-Dieter Arndt, Peter Bieling, and Stefan Rausner. “Structural transitions of F-actin upon ATP hydrolysis at near-atomic resolution revealed by cryo-EM”. In: Nature Structural & Molecular Biology 25 (2018), pp. 528–537.

51. HP Erickson. “Co-operativity in protein-protein association. The structure and stability of the actin filament”. In: Journal of molecular biology 206.3 (1989), pp. 465–474.

52. A Bremer, R C Millonig, R Sütterlin, A Engel, T D Pollard, and U Aebi. “The structural basis for the intrinsic disorder of the actin filament: the “lateral slipping” model.” In: The Journal of Cell Biology 115.3 (1991), pp. 689–703.

53. Jhih-Wei Chu and Gregory A. Voth. “Allostery of actin filaments: Molecular dynamics simulations and coarse-grained analysis”. In: PNAS 201 (2005), pp. 13111–13116.

54. Mark A. Rould, Qun Wan, Peteranne B. Joel, Susan Lowey, and Kathleen M. Trybus. “Crystal structures of expressed non-polymerizable monomeric actin in the ADP and ATP states”. In: Journal of Biological Chemistry 281 (2006), pp. 31909–31919.

55. Zeynep A. Oztug Durer, Dmitri S. Kudryashov, Michael R. Sawaya, Christian Altenbach, Wayne Hubbell, and Emil Reisler. “Structural States and Dynamics of the D-Loop in Actin”. In: Biophysical Journal 103.5 (2012), pp. 930–939.

56. Mahito Kikumoto and Fumio Oosawa. “Thermodynamic measurements of actin polymerization with various cation species”. In: Cytoskeleton 74.12 (2017), pp. 465–471.

57. Panagiotis L. Kastritis, João P.G.L.M. Rodrigues, Gert E. Folkers, Rolf Boelens, and Alexandre M.J.J. Bonvin. “Proteins Feel More Than They See : Fine-Tuning of Binding Affinity by Properties of the Non-Interacting Surface”. In: Journal of Molecular Biology 426 (2014), pp. 2632–2652.

58. Xiaorui Chen, Fengyun Ni, Xia Tian, Elena Kondrashkina, Qinghua Wang, and Jianpeng Ma. “Structural Basis of Actin Filament Nucleation by Tandem W Domains”. In: Cell Reports 3.6 (2013), pp. 1910–1920.

59. Peteranne B. Joel, Patricia M. Fagnant, and Kathleen M. Trybus. “Expression of a Nonpolymerizable Actin Mutant in Sf9 Cells”. In: Biochemistry 43.36 (2004), pp. 11554–11559. issn: 0006-2960.

60. Mark A. Rould, Qun Wan, Peteranne B. Joel, Susan Lowey, and Kathleen M. Trybus. “Crystal Structures of Expressed Non-polymerizable Monomeric Actin in the ADP and ATP States”. In: Journal of Biological Chemistry 281.42 (2006), pp. 31909–31919.

61. Anna M. Ducka, Peteranne Joel, Grzegorz M. Popowicz, Kathleen M. Trybus, Michael Schleicher, Angelika A. Noegel, Robert Huber, Tad A. Holak, and Tomasz Sitar. “Structures of actin-bound Wiskott-Aldrich syndrome protein homology 2 (WH2) domains of Spire and the implication for filament nucleation”. In: Proceedings of the National Academy of Sciences 107.26 (2010), pp. 11757–11762.

62. Weihua Zheng, Nicholas P. Schafer, Aram Davtyan, Garegin A. Papoian, and Peter G. Wolynes. “Predictive energy landscapes for protein–protein association”. In: Proceedings of the National Academy of Sciences 109.47 (2012), pp. 19244– 19249.

63. Ilya A. Vakser. “Protein-Protein Docking: From Interaction to Interactome”. In: Biophysical Journal 107.8 (2014), pp. 1785–1793.

64. Elena E. Grintsevich, Martin Phillips, Dmitry Pavlov, Mai Phan, Emil Reisler, and Andras Muhlrad. “Antiparallel Dimer and Actin Assembly”. In: Biochemistry 49.18 (2010), pp. 3919–3927.

65. Naoki Watanabe. “Inside view of cell locomotion through single-molecule: fast F-/G-actin cycle and G-actin regulation of polymer restoration.” In: Proc Jpn Acad Ser B Phys Biol Sci 86.1 (2010), pp. 62–83.

66. Otto G. Berg and Peter H. von Hippel. “Diffusion-Controlled Macromolecular Interactions”. In: Annual Review of Biophysics and Biophysical Chemistry 14.1 (1985), pp. 131–158.

67. S H Northrup and H P Erickson. “Kinetics of protein-protein association explained by Brownian dynamics computer simulation.” In: Proceedings of the National Academy of Sciences 89.8 (1992), pp. 3338–3342.

68. Ludovic R. Otterbein, Philip Graceffa, and Roberto Dominguez. “The Crystal Structure of Uncomplexed Actin in the ADP State”. In: Science 293.5530 (2001), pp. 708–711.

69. Xiange Zheng, Karthikeyan Diraviyam, and David Sept. “Nucleotide Effects on the Structure and Dynamics of Actin”. In: Biophysical Journal 93.4 (2007), pp. 1277–1283.

70. I. Fujiwara, D. Vavylonis, and T. D. Pollard. “Polymerization kinetics of ADP- and ADP-Pi-actin determined by fluorescence microscopy”. In: Proc. Natl. Acad. Sci. U. S. A. 104 (2007), pp. 8827–8832.

71. Chun Tang, Junji Iwahara, and G. Marius Clore. “Visualization of transient encounter complexes in protein-protein association”. In: Nature 444.7117 (2006), pp. 383–386.

72. R. Killick, P. Fearnhead, and A. Eckley. “Optimal Detection of Changepoints With a Linear Computational Cost”. In: Journal of the American Statistical Association 107 (2012), pp. 1590–1598.

73. Nancy R. Zhang and David O. Siegmund. “A Modified Bayes Information Criterion with Applications to the Analysis of Comparative Genomic Hybridization Data”. In: Biometrics 63.1 (2007), pp. 22–32.

