## Supplemental Text and Figures for "Insights into actin polymerization and nucleation using a coarse grained model"

##### Additional information on simulation methods

The interaction potential between amino acids $i$ & $j$ at distance $r$ are defined by the KH model as:

$$u_{ij}(r) = \begin{cases} 4|\epsilon_{ij}|[(\frac{\sigma_{ij}}{r})^{12} - (\frac{\sigma_{ij}}{r})^{6}] \,, & \epsilon_{ij} \leq 0 \\ 4\epsilon_{ij}[(\frac{\sigma_{ij}}{r})^{12} - (\frac{\sigma_{ij}}{r})^{6}] + 2\epsilon_{ij} \,, & r \leq 2^{\frac{1}{6}}\sigma_{ij},\ \epsilon_{ij} \geq 0 \\ -4\epsilon_{ij}[(\frac{\sigma_{ij}}{r})^{12} - (\frac{\sigma_{ij}}{r})^{6}] \,, & r \geq 2^{\frac{1}{6}}\sigma_{ij},\ \epsilon_{ij} \geq 0 \end{cases}$$

This potential was implemented in LAMMPS through modification of the source code [1]. The $\epsilon_{ij}$ values are obtained from:

$$\epsilon_{ij} = \lambda(e_{ij} - e_0)$$

For KH Model A used in this paper, $\lambda = 0.159$ and $e_0 = -2.27\, k_B\, T_0$, where $T_0 = 298\, K$ was a fixed number as $T$ was varied in the simulations. The value for $e_{ij}$ is the corresponding value in the Miyazawa-Jernigan pairwise interaction matrix, table 3 in reference [2]. The interaction radii, $\sigma_{ij}$, are as defined in equation 6 and table 5 of reference [3]. In the KH model, residue charges for electrostatic interactions correspond to pH 7 such that the charge is $+e$ for Lys and Arg, $-e$ for Asp and Glu, and $+0.5e$ for His, where $e$ is the elementary charge.

Simulations in LAMMPS were performed with a timestep of 10 fs. Typical simulations are of the order of microseconds to 10s of microseconds, with specific convergence criteria are discussed in the main text. For all MD cases we used a temperature list with 16 replicas between 180 K and 300 K, where temperature was maintained with the Langevin dynamics thermostat, using the fix_langevin command in LAMMPS with a damping parameter of 1000 fs. The temperature list was tuned for sufficiently high acceptance rates and well-distributed fraction of each replica in each temperature. To ensure sufficient replica sampling, swaps are attempted every 1000 time steps. Typical swap success rates were in the range 30-80%. The final temperature list in degrees Kelvin used was 180, 182, 185, 188, 191, 194, 197, 201, 205, 209, 214, 220, 240, 260, 280, & 300. Additionally, unless otherwise stated, all simulations were performed at 4 different concentrations, corresponding to cubic box side lengths of 368 Å, 464 Å, 629 Å, and 793 Å.

Unless otherwise stated, all proteins and protein assemblies were treated as single rigid bodies by way of the fix_rigid command in LAMMPS, which sums all forces and torques on the individual particles of the rigid body and causes the unit to move together as one body, rather than as individual constituents. Masses of rigid bodies were reduced to decrease convergence time [1]. Sample LAMMPS input files are provided for reference as supplemental material.

##### Contacts formed between subunits

In this section we provide comments related to Fig. 3C showing the contacts maps of incoming F-Oda monomer with F-Oda dimer. As expected for BE binding, contacts along the long-pitch helix direction, between incoming subunit $a$ and subunit $c$, involve the D-loop (residues 37-47) of the incoming subunit. In the BE1 structure, these residues made the strongest contacts with residues around the hydrophobic groove of subunit $c$ (residues 165-171, 285-289, 351-355, and 373-375, including contacts with C374 [4]). Further residues in subdomain 2 (61-66) of the incoming subunit form contacts with subdomain 3 (166-169 and 286-289) of subunit $c$. Additional long-pitch contacts include residues in subdomain 4 of the incoming subunit (198-208 and 240-248) making contacts with subdomain 3 of subunit $c$ (in 285-289 and 287-294/322-326 regions, respectively).

Short-pitch helix contacts for BE1 binding between incoming subunit $a$ and subunit $b$ include residues in subdomain 2 (64-80) of the incoming subunit that form contacts with residues of subdomain 3 (173-174, 269-272, 276-290, and 319-322) of subunit *b*. Further, residues 36-41 of the D-loop also make short-pitch helix contacts with subdomain 4 of of subunit *b* (residues 264-272). Residues in subdomain 1 (109-115) form contacts with residues with subdomain 3 (283-284 and 286-290) of subunit *b*. Further, residues in subdomain 3 (176-181) and 4 (183-204) of the incoming subunit form contacts with residues in subdomain 1 (75, 109-118) and subdomain 3 (171-179, 269-271, 284 & 375) of subunit *b*. Residues in subdomain 4 (263-271) of the incoming subunit form contacts with residues subdomain 4 (110, 113) and subdomain 3 (168-175, 370-375) of subunit *b*. Residues 158-160 (near the nucleotide binding cleft) of the incoming subunit also form contacts with residues in subdomain 3 (173, 284-287) of subunit *b*. Residue S14, also near the nucleotide binding cleft, of the incoming subunit forms strong contacts with K284 (in subdomain 3) of subunit b.

PE1 binding is essentially symmetric to BE1 binding but the subunit is at the other end. The remaining hotspots in Fig. 3C correspond to the other complexes (BE2, PE2, AP1, AP2).

| State | $K_d$ $(\mu M)$ |
|---|---|
| F-Oda dimer + F-Oda monomer (4$\sigma$, 300 K) | |
| BE1+BE2 | 440 |
| PE1+PE2 | 163.5 |
| AP1 | 110 |
| AP2 | 2800 |
| F-Oda dimer + F-Oda monomer (3$\sigma$, 197 K) | |
| BE1+BE2 | 0.15 $\pm$ 0.04 |
| PE1+PE2 | 0.13 $\pm$ 0.03 |
| AP1 | 0.31 $\pm$ 0.19 |
| AP2 | 9.0 $\pm$ 0.9 |
| F-Oda flex. dimer + G-ATP-Grac. flex. monomer (3$\sigma$, 197 K) | |
| BE1+BE2+BE3 | 0.035 $\pm$ 0.009 |
| AP1 | 0.51 $\pm$ 0.07 |
| AP2 | 0.08$\pm$ 0.03 |
| O1+O2+O3+04 | 0.015$\pm$ 0.006 |

Table S1: Binding affinities for monomer binding to fixed dimer, calculated using the KH model as described in the main text. The table indicates the values for KH Model A with cutoff $3\sigma$ or $4\sigma$ and simulated temperature.

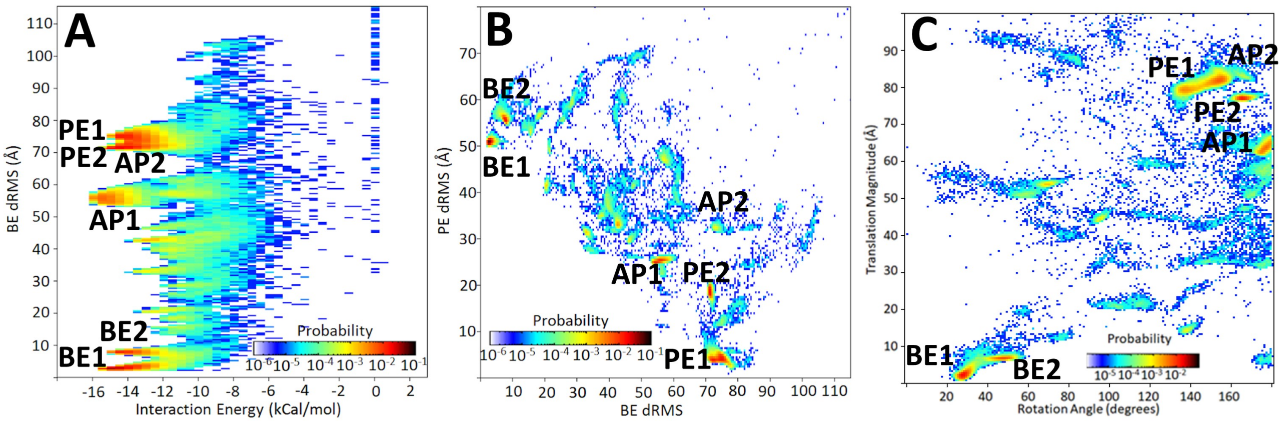

Figure S1: Representation of ensemble of bound states for simulations of Figs. 1 and 2. (A) Interaction energy of versus dRMS distance of incoming monomer to the BE reference structure by Oda et al. [5]. All high probability states have similar energy. Very high dRMS can lead to degeneracy in regions, which is why the PE states overlap in this space. (B) Plot showing dRMS distance of incoming monomer to both BE and PE reference structure by Oda et al. This space helps distinguish among states and is used to cluster the data. (C) Same as Fig. 2A. This space describes the relative translation and rotation of the monomer in the simulation with respect to the BE state in the Oda et al.

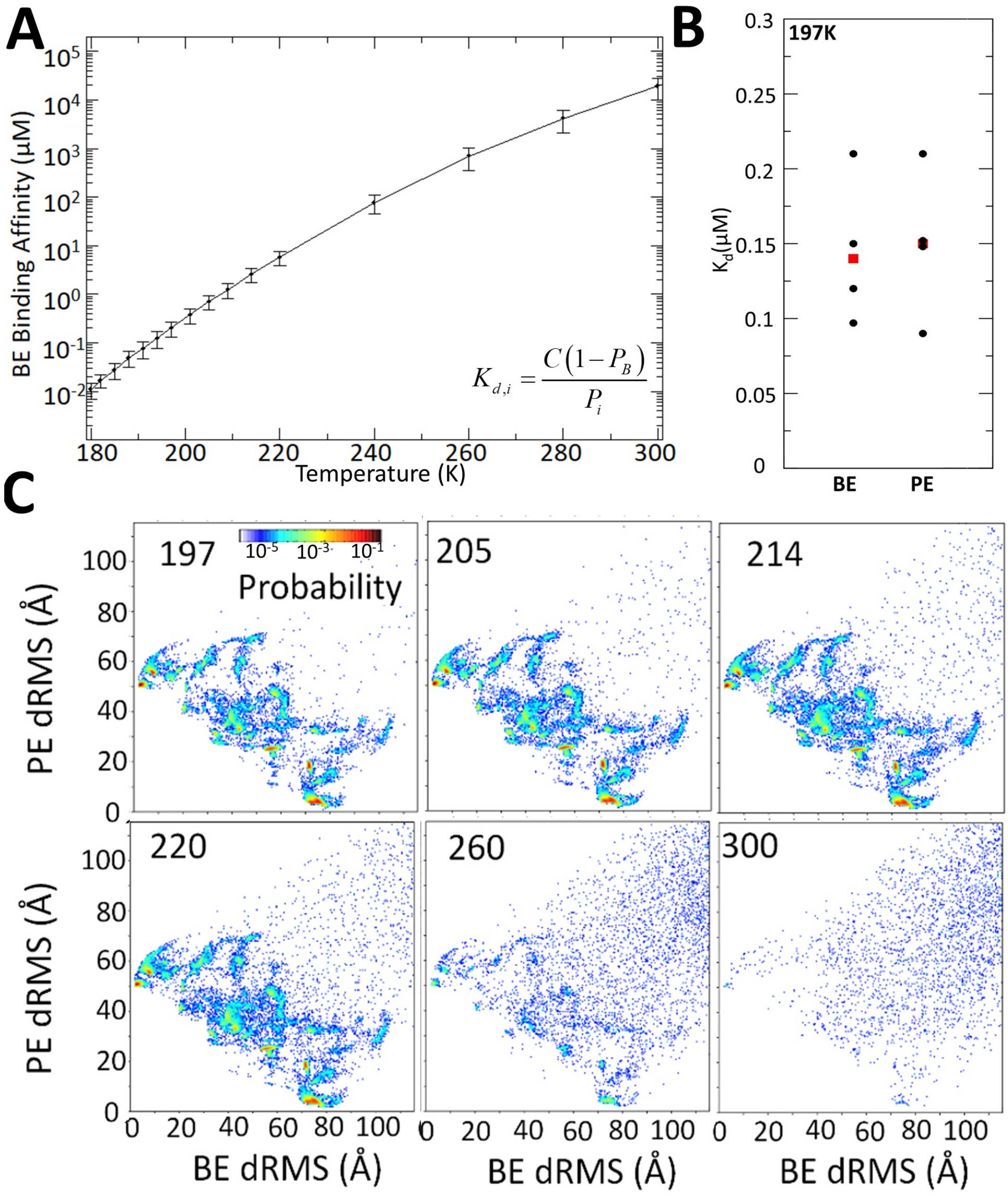

Figure S2: Binding affinity and effect of simulation temperature. (A) BE1 binding affinity of F-Oda rigid dimer interacting with rigid F-Oda monomer as a function of temperature. Other parameters same as Fig. 1. Error bar shows standard deviation calculated from four separate simulations. (B) Dissociation constant for BE and PE at 197K. Black shows calculation from four simulations and red shows average. Binding affinity at BE and PE are similar. In this plot the BE and PE states are defined as those frames that have dRMS ¡ 10 Å to the corresponding structure in the model by Oda et al. [5] (they include BE1 and BE2 for the BE and PE1 for the PE, see Fig. 1). (C) Two-dimensional dRMS plots as Fig. S1B show the hotspots remain the same as function of temperature but sampled with lower probabilities with increasing temperature.

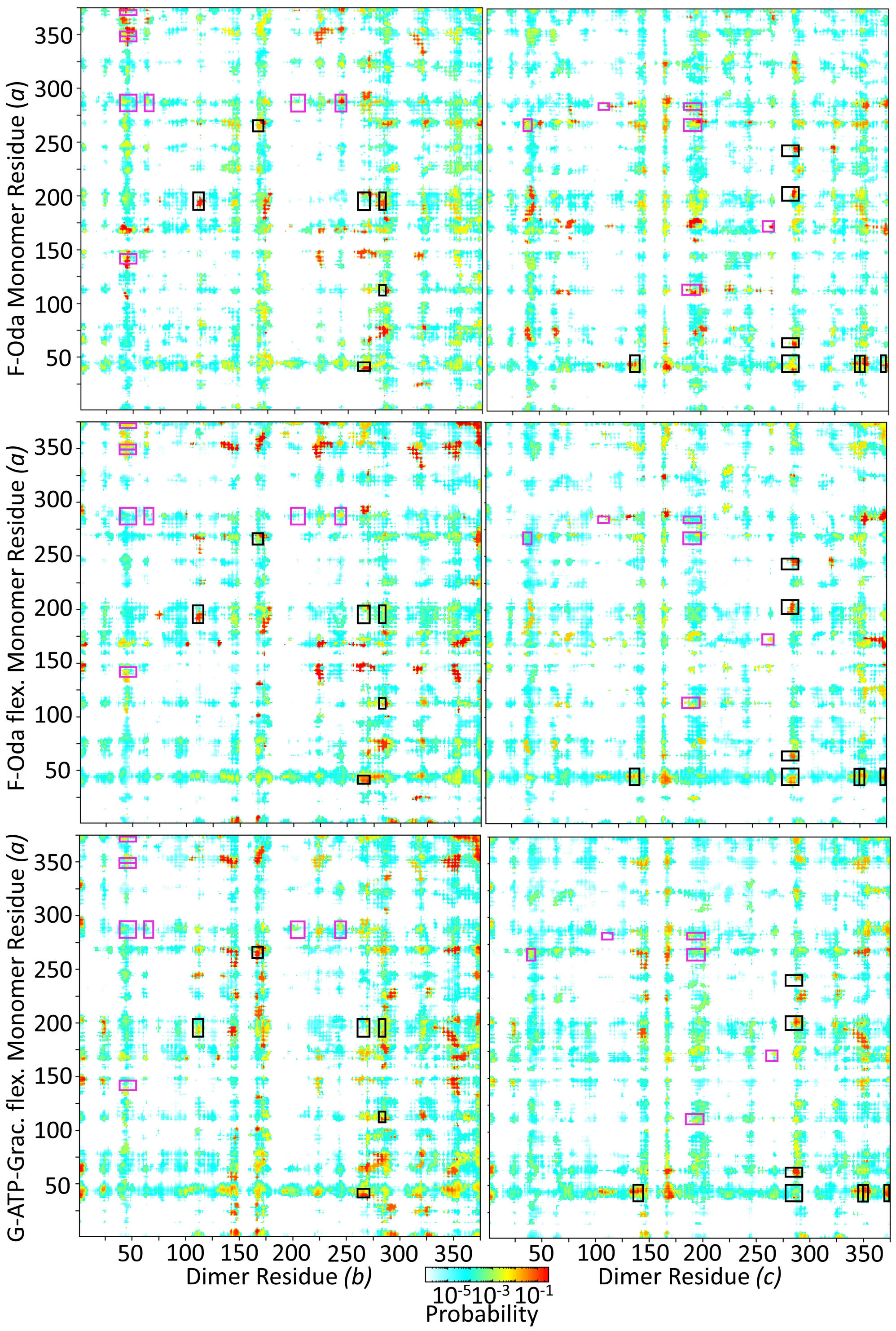

Figure S3: Contact maps for incoming monomer (subunit *a*) associating with fixed dimer (subunits *b* and *c*) corresponding to the simulations of Fig. 2, 4 and 5 where the incomoning monomer was rigid F-Oda (top row, same as Fig. 3C), F-Oda flex (middle row), and G-ATP-Grac. flex (bottom row), respectively.

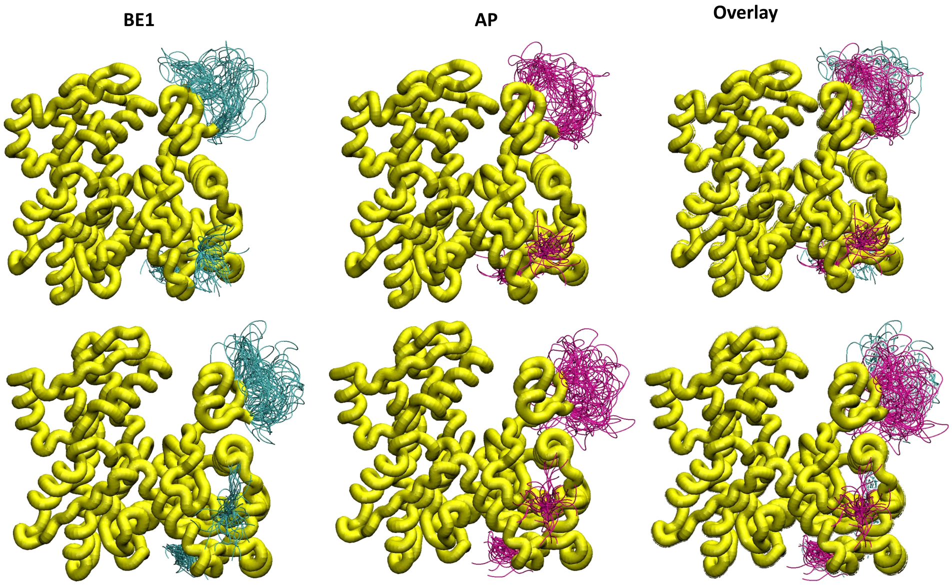

Figure S4: Images comparing the typical bound and unbound conformations of the flexible regions in the F-Oda flex monomer simulations (Fig. 4). Roughly, when the system is in BE1, the D-loop is associated with the dimer and the termini are unbound; when the system is in AP, the D-loop is free and the termini are associated to the dimer. These images show that the D-loop remains flexible in all cases, adopting a slightly more elongating configuration in the BE1 state.

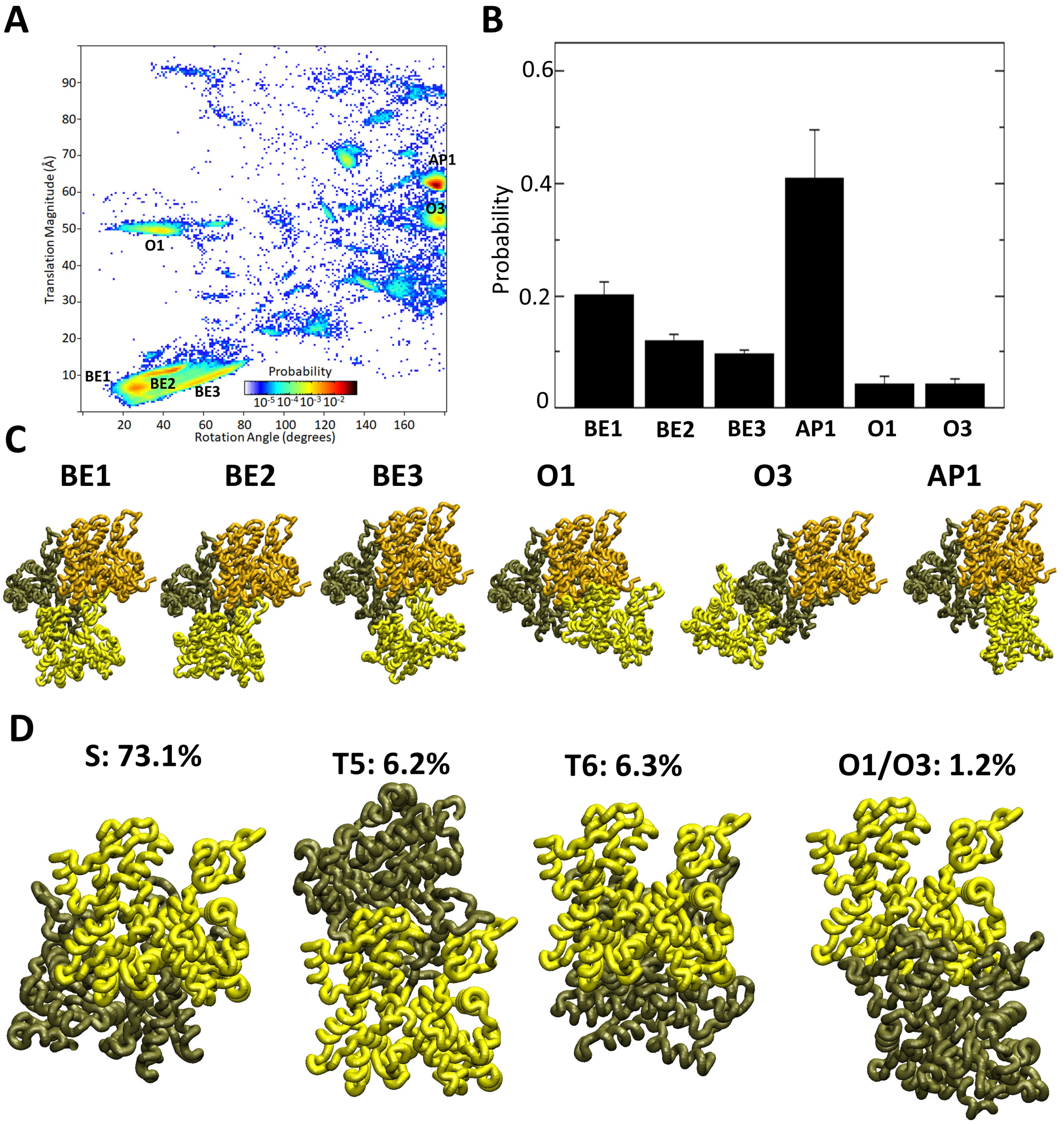

Figure S5: Simulation with G-ATP-Grac. rigid monomer. (A) Same as in Figure 2, but with the subunit as a G-actin monomer. (B) Probabilities of the different complexes identified in A. (C) Images of the high probability complexes. (D) Images of high probability complexes from monomer-monomer simulations with two G-ATP-Grac. rigid monomers. Of the short-pitch, long-pitch, and antiparallel dimer complexes, only the short-pitch was identified. The parameters for panels A-C are the same as for Figs. 2,4,5. For panel D, simulation parameters are the same as Fig. 6 except the box size is 300 Å.

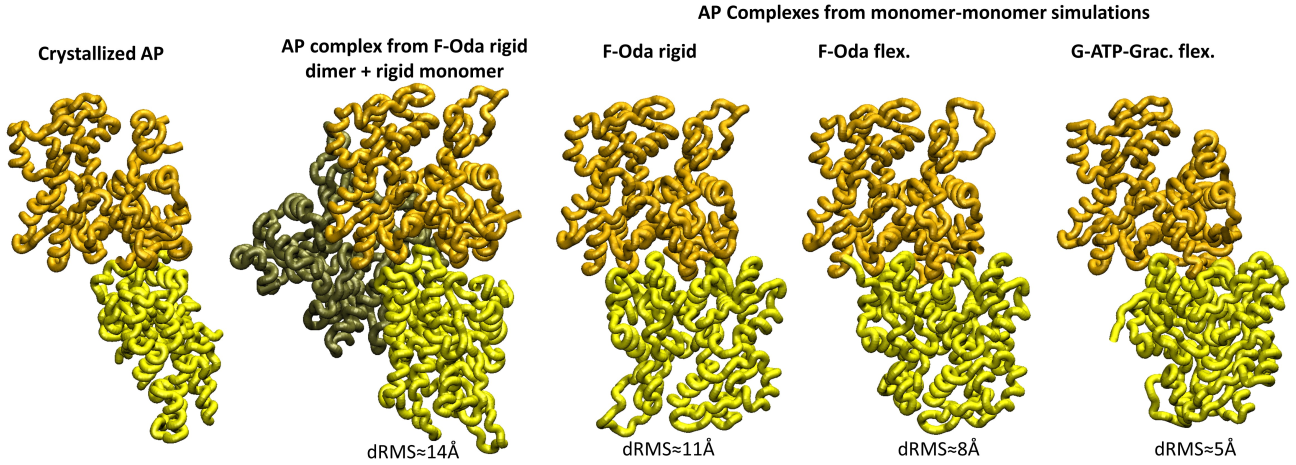

Figure S6: Simulation of AP complexes and comparison to the antiparallel dimer crystal structure 1RFQ in [6]. dRMS of simulation complexes in the AP region as defined in Fig. 2 with respect to 1RFQ.

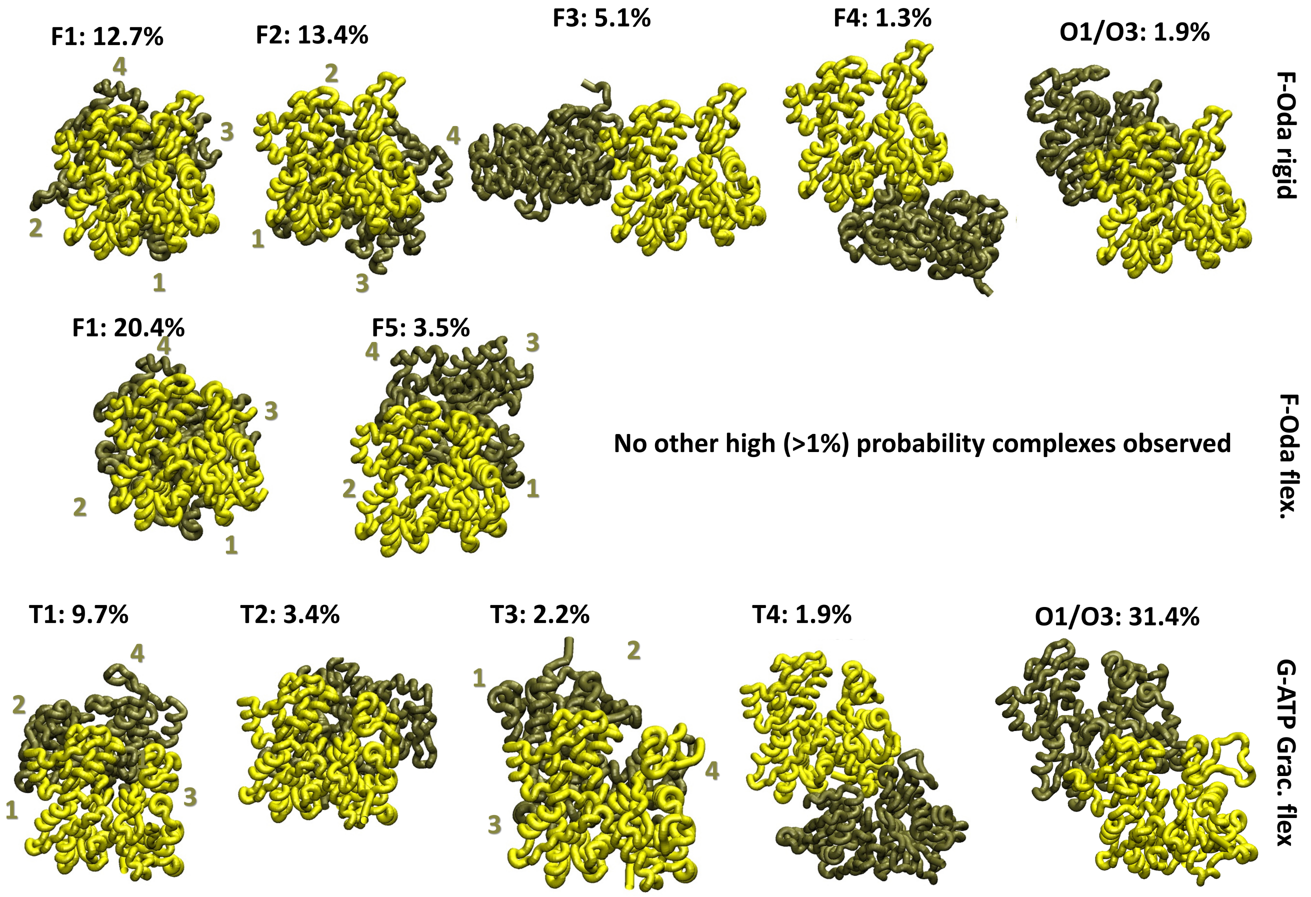

Figure S7: Additional complexes identified in the monomer-monomer simulations of Fig. 6. The subdomains of some actin monomers are numbered.

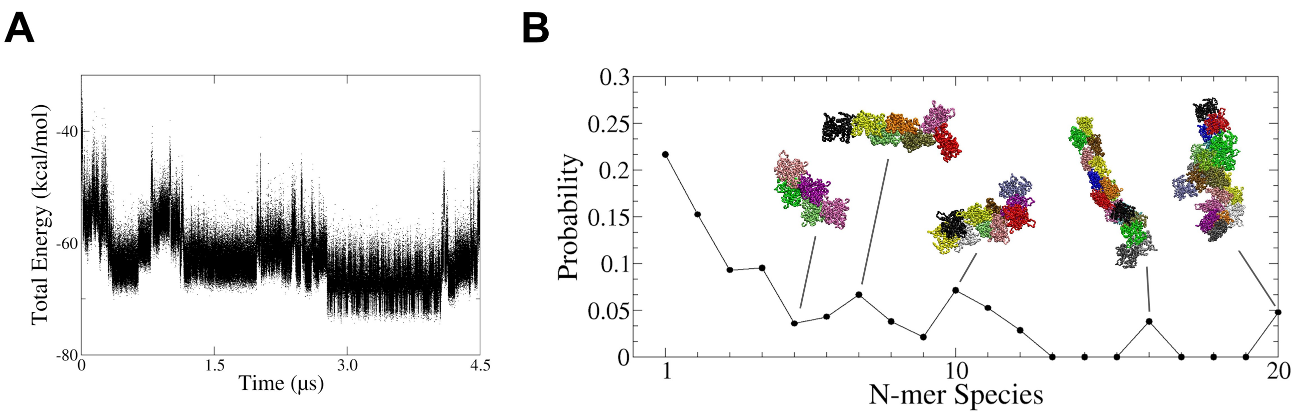

Figure S8: Quantification of results in Fig. 7. (A) Potential energy versus time of simulation in Fig. 7A (6-monomer long filament in the Oda et al. [5] configuration) shows stability of filament configuration. (B) Summary of end state of $n = 21$ simulations run with parameters listed for Fig. 7B (serial MD simulations of a pool of 20 rigid F-Oda monomers in a box at 204 K after 0.94-2.8 $\mu s$). Probability for an individual monomer to be found in a given N-mer species is shown. A subunit is considered to be part of an N-mer by visually checking monomers in contact. Snapshots show examples of N-mers of indicated size.

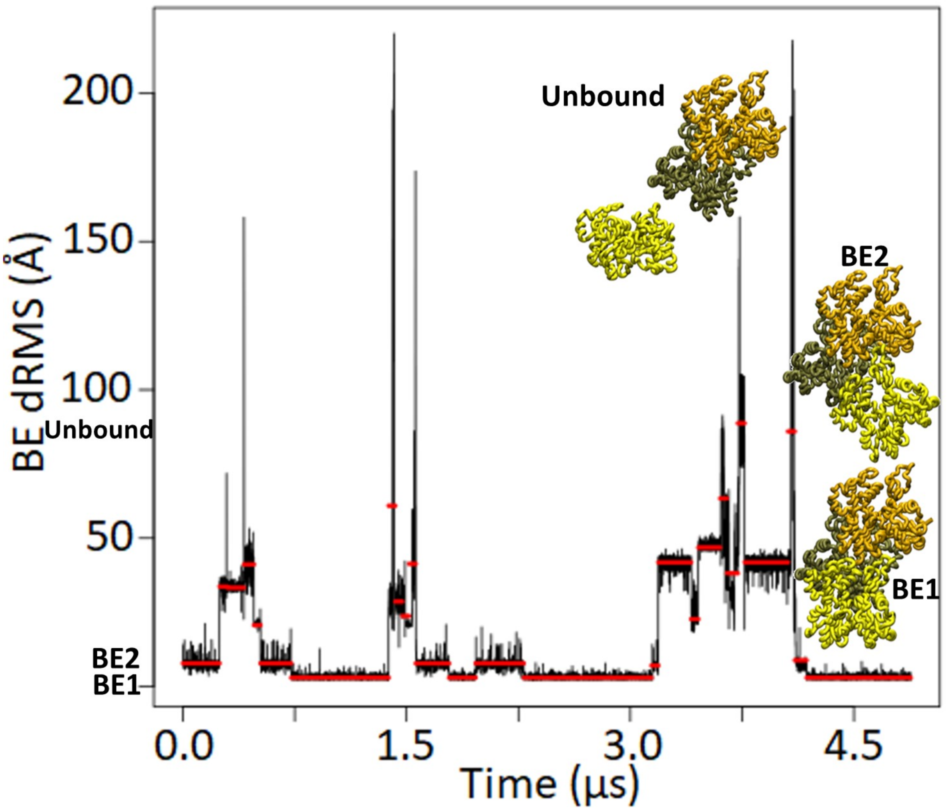

Figure S9: Serial MD simulations of the rigid F-Oda dimer plus rigid F-Oda monomer. Multiple transitions between different states were observed. Red lines identify parts of the trajectory where the mean is roughly the same, as identified by change-point analysis using the PELT algorithm [7] with an MBIC penalty [8]. We found that the BE2 may be an intermediate in association from unbound to BE1.

### Movies

Movie S1. Serial MD simulation showing typical states of F-Oda rigid dimer interacting with a rigid F-Oda monomer, including a BE association event, T=197K. Snapshots have been aligned such that the diffusing fixed dimer remains in the center. The incoming monomer starts in the AP1 state, later transitions to BE1, alternates between BE1/2, then dissociates and binds to the PE as PE1, and finally dissociates and rebinds to BE1/2.

Movie S2: Images of AP1 states from various models are compared.

Movie S3. Serial MD simulation showing typical states of F-Oda flex. dimer interacting with G-actin (G-ATP-Grac. flex.), T=197K. Snapshots have been aligned such that the diffusing fixed dimer remains in the center. The incoming monomer starts in AP1, transitions to the barbed end exploring states BE1,BE2 and BE3 and finally transitions to O3.

Movie S4. Serial MD simulation showing F-Oda flex. dimer barbed end association with a G-ATP-Grac. flex monomer. Same as Movie S2 but with a different sequence of events showing a case where the D-loop latches on and guides incoming monomer to rotate into place to the BE2/3 states.

Movie S5: Serial MD simulation showing F-Oda flex. dimer barbed end association with a G-ATP-Grac. flex monomer. Same as Movie S2 but with a different sequence of events showing a case where the subdomain 4 of the incoming monomer binds first, followed by the D-loop, and then subunit rotates into place to the BE2/3.

#### References

[1] Brandon G. Horan et al. "Computational modeling highlights the role of the disordered Formin Homology 1 domain in profilin-actin transfer". In: FEBS Lett. 592 (2018), pp. 1804–1816.

[2] Sanzo Miyazawa and Robert L. Jernigan. "Residue – Residue Potentials with a Favorable Contact Pair Term and an Unfavorable High Packing Density Term, for Simulation and Threading". In: Journal of Molecular Biology 256.3 (1996), pp. 623–644.

[3] Gerhard Hummer and Young Kim. "Coarse-grained Models for Simulations of Multiprotein Complexes: Application to Ubiquitin Binding". In: Journal of Molecular Biology 375 (2008), pp. 1416–1433.

[4] Steven Z. Chou and Thomas D. Pollard. "Mechanism of actin polymerization revealed by cryo-EM structures of actin filaments with three different bound nucleotides". In: Proceedings of the National Academy of Sciences 116.10 (2019), pp. 4265–4274.

[5] Toshiro Oda et al. "The nature of globular- to fibrous-actin transition". In: Nature 457 (2009), pp. 441–445.

[6] Robbie Reutzel et al. "Actin crystal dynamics: structural implications for F-actin nucleation, polymerization, and branching mediated by the anti-parallel dimer". In: Journal of Structural Biology 146 (2004), pp. 291–301.

[7] R. Killick, P. Fearnhead, and A. Eckley. "Optimal Detection of Changepoints With a Linear Computational Cost". In: Journal of the American Statistical Association 107 (2012), pp. 1590–1598.

[8] Nancy R. Zhang and David O. Siegmund. "A Modified Bayes Information Criterion with Applications to the Analysis of Comparative Genomic Hybridization Data". In: Biometrics 63.1 (2007), pp. 22–32.
